# Mitophagy at the oocyte-to-zygote transition promotes species immortality

**DOI:** 10.1101/2025.02.01.636045

**Authors:** Siddharthan Balachandar Thendral, Sasha Bacot, Katherine S. Morton, Qiuyi Chi, Isabel W. Kenny-Ganzert, Joel N. Meyer, David R. Sherwood

## Abstract

The quality of inherited mitochondria determines embryonic viability^1^, metabolic health during adulthood and future generation endurance. The oocyte is the source of all zygotic mitochondria^2^, and mitochondrial health is under strict developmental regulation during early oogenesis^3–5^. Yet, fully developed oocytes exhibit the presence of deleterious mitochondrial DNA (mtDNA)^6,7^ and mitochondrial dysfunction from high levels of endogenous reactive oxygen species^8^ and exogenous toxicants^9^. How fully developed oocytes prevent transmission of damaged mitochondria to the zygotes is unknown. Here we discover that the onset of oocyte-to-zygote transition (OZT) developmentally triggers a robust and rapid mitophagy event that we term mitophagy at OZT (MOZT). We show that MOZT requires mitochondrial fragmentation, activation of the macroautophagy system and the mitophagy receptor FUNDC1, but not the prevalent mitophagy factors PINK1 and BNIP3. Oocytes upregulate expression of FUNDC1 in response to diverse mitochondrial insults, including mtDNA mutations and damage, uncoupling stress, and mitochondrial dysfunction, thereby promoting selection against damaged mitochondria. Loss of MOZT leads to increased inheritance of deleterious mtDNA and impaired bioenergetic health in the progeny, resulting in diminished embryonic viability and the extinction of descendent populations. Our findings reveal FUNDC1-mediated MOZT as a mechanism that preserves mitochondrial health during the mother-to-offspring transmission and promotes species continuity. These results may explain how mature oocytes from many species harboring mutant mtDNA give rise to healthy embryos with reduced deleterious mtDNA.

## Main

Mitochondria are universal in their role as energy production and signaling centers that sustain eukaryotic life. Arising through a pivotal endosymbiosis event ∼1.5 billion years ago, mitochondria still maintain independent genomes (mtDNA) that encode proteins essential for energy production^10^. Mitochondria are inherited maternally in most eukaryotes and inheritance of functional mitochondria is crucial to support energetic, signaling, and metabolic demands of embryonic development^1,2^. Importantly, late oocytes use and pass along the founding copies of mtDNA that serve as a template for expansion throughout offspring life.

Mitochondrial quality is fundamental to organismal and population health. Deleterious mtDNA and subsequent mitochondrial dysfunction are associated with embryonic lethality, metabolic disorders, cancer, neurodegeneration, and infertility^11,12^. Furthermore, mtDNA mutations, when accumulated over generations are irreversible and could lead to species extinction^13,14^. Many factors disrupt oocyte mitochondrial and mtDNA health, such as maternal diet, toxicants, radiation, reactive oxygen species and mtDNA replication errors^9,15^. Since mitochondria lack robust DNA repair machinery^16^, mtDNA is more prone to accumulation of damage than nuclear DNA, and animal mtDNA mutates more quickly over evolutionary time than nuclear DNA^17^. Yet, despite the high mtDNA mutation rate, few deleterious mtDNA variants transmit over generations, indicating that there is selection against deleterious mtDNA^18^.

Oocyte maturation is energetically demanding and requires a critical threshold of mitochondrial abundance^8,19^. Deficits in mtDNA and mitochondrial derived ATP in oocytes leads to fertilization failure and impaired embryonic development^20^. This poses a challenge for the mature oocyte - maintaining abundant, highly active mitochondria but then ensuring transfer of high-quality mitochondria to the zygote. While oocyte mitochondrial quality control has been extensively studied in the context of mitochondrial bottlenecking^21,22^ and programmed cell death^23,24^, little is known about whether the oocyte counteracts damaged mitochondria.

Suggestive of organelle level purifying selection, mutant mtDNA variants are common in mature oocytes from many species^6,25–27^, but are lost in the resulting progeny. Notably, a 4977-bp deletion called the “common deletion” has been detected in 30-50% of human oocytes, along with several mtDNA rearrangements. Intriguingly, the frequency of such deleterious mtDNA diminishes ∼2-fold in embryos generated from these oocytes^7,28^. Similarly, mice oocytes harboring abundant deleterious mtDNA give rise to progeny free of mutations^6^. In *C. elegans*, mutant mtDNA variants are also reduced during transmission from mother to progeny^23^.

Together, these studies support selection against deleterious mtDNA at the mitochondrial level within the oocyte during transmission to offspring. However, the mechanisms underlying mitochondrial purifying selection in the oocyte, and the impacts of loss of selection have remained elusive^29^.

Here, we uncover a rapid programmed mitochondrial selection event in the *C. elegans* oocyte that specifically occurs during the oocyte-to-zygote transition (OZT), which we call mitophagy at OZT (MOZT). We identify FUNDC1, a conserved mitochondrial membrane protein^30^, as the molecular link that coordinates mitochondrial fragmentation and mitophagy during OZT and show that loss of FUNDC1 increases the frequency of inherited mutant mtDNA, which leads to mutational meltdown and population extinction over multiple generations.

## Results

### Programmed mitochondrial reduction at OZT

*C. elegans* oogenesis begins with syncytial germline stem cells proliferating in the mitotic zone located at each distal end of a u-shaped germline. Germ cells exit the mitotic zone, initiate differentiation and enter meiotic prophase I. They then move along the germline proximally while going through pachytene and diplotene stages. As the germ cells progress along the germline bend, they complete cellularization, increase in cellular and nuclear volume and finally become oocytes, arresting in diakinesis of meiotic prophase I^31^ (Fig. 1a). As more germ cells undergo this process, newly formed oocytes stack at the proximal end of the gonad. The oldest and most proximal oocyte is referred to as the −1 oocyte, followed distally by younger −2, −3 oocytes, etc. (Fig. 1a). While the −2, −3 and other oocytes remain arrested in meiotic prophase I, major sperm protein (MSP) released from sperm triggers the −1 oocyte to undergo the oocyte-to-zygote transition (OZT). OZT occurs over a rapid ∼20 min time period and involves many cellular processes, such as nuclear envelope breakdown, cortex rearrangement, ovulation, and finally fertilization by sperm to become the zygote^32^ (Fig. 1a).

**Fig.1:**
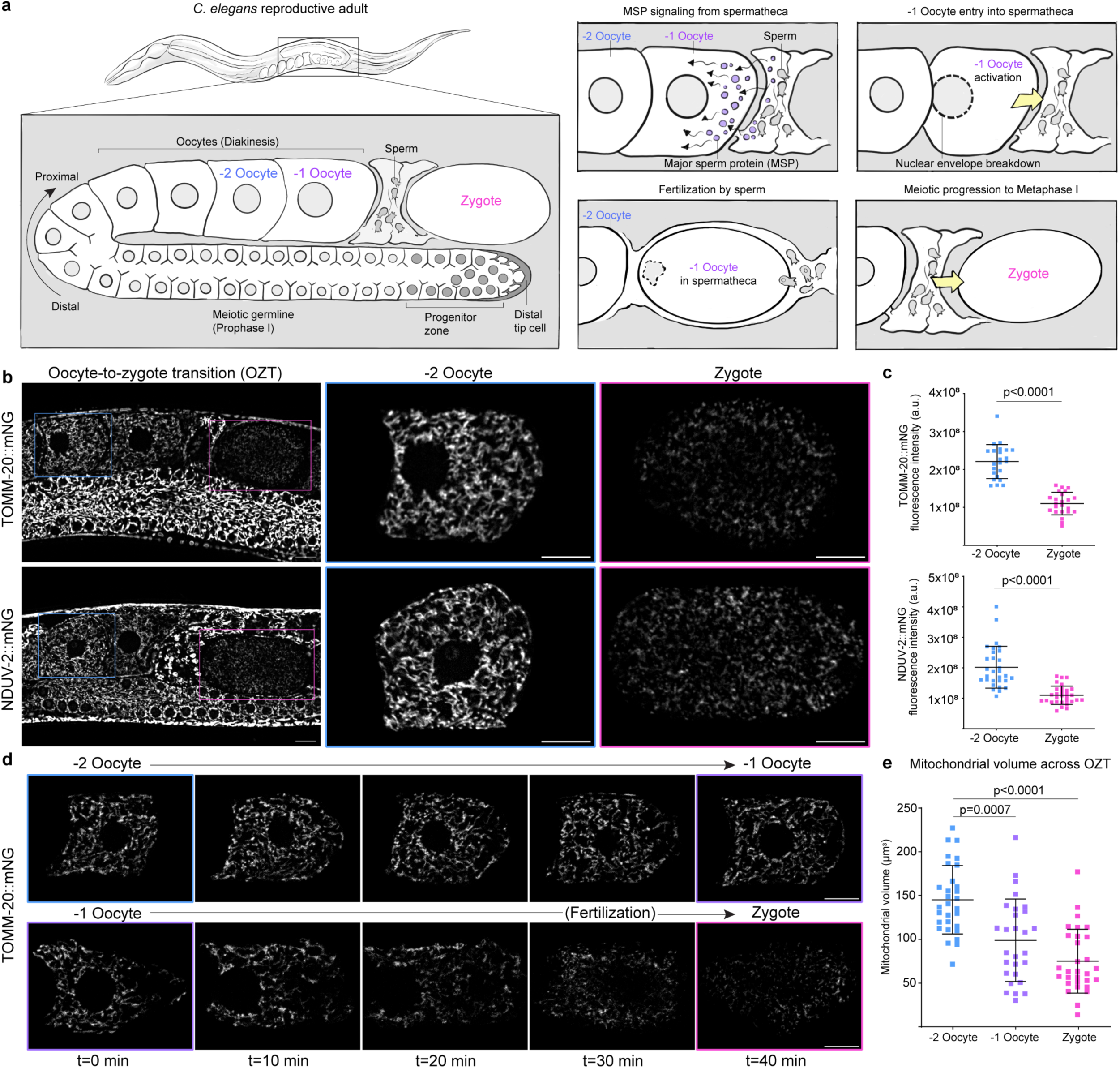
Mitochondrial reduction during the *C. elegans* oocyte-to-zygote transition. **a**, A schematic of the adult *C. elegans* hermaphrodite germline (left) and a timeline of the oocyte-to-zygote transition (OZT) (right). **b**, Confocal fluorescence images of mitochondria during *C. elegans* OZT. Endogenous outer mitochondrial membrane protein TOMM-20::mNG (top) and inner mitochondrial membrane protein NDUV-2::mNG (bottom) are reduced in zygotes compared to −2 oocytes. Blue boxes indicate −2 oocytes and pink boxes zygotes. The −2 oocytes and zygotes enclosed in boxed regions are magnified. Scale bars, 10µm. **c**, Total fluorescence intensity of endogenous TOMM-20::mNG (top) and NDUV-2::mNG (bottom) in −2 oocytes and zygotes. The data represent the mean ± s.d. (n = 30 animals per strain) from three biological replicates. P values were calculated using two-tailed Student’s *t*-tests. **d**, Time-lapse images showing changes in mitochondrial abundance over 40 min in −2 and −1 oocytes expressing endogenous TOMM-20::mNG (*tomm-20::mNG*). Fluorescence images show unchanged mitochondrial abundance in −2 oocyte throughout its progression to the −1 position (top), compared to mitochondrial reduction in −1 oocyte during transition to zygote (bottom). Data is representative of six independent experiments. **e**, Mitochondrial volume (µm³) determined from endogenous NDUV-2 (*nduv-2::mNG*) signal distribution in −2 oocytes, −1 oocytes and zygotes. The data represent the mean ± s.d. (n = 30 animals) from three biological replicates. P values were calculated using Kruskal Wallis followed by Dunn’s multiple comparisons test.

To visualize mitochondria during oocyte development, we used genome editing to knock-in mNeonGreen (mNG) into the outer mitochondrial membrane (OMM) protein TOMM-20::mNG and inner mitochondrial membrane (IMM) protein NDUV-2::mNG. Strikingly, live imaging revealed a ∼2-fold reduction in mitochondrial proteins (Fig. 1b,c) and mitochondrial volume (Fig. 1e) from the −2 oocyte to zygote. Labelling mitochondria using the general mitochondrial dye nonyl acridine orange (NAO) confirmed a similar decrease in mitochondrial content (Extended Data Fig. 1a,b). Live-cell imaging of mitochondria in arrested oocytes (−2 oocyte) and the oocyte undergoing OZT (−1 oocyte) revealed the reduction in mitochondrial volume was specific to −1 oocytes (Fig. 1d and Extended Data Fig. 1c-f). Mitochondrial reduction initiated with the onset of OZT in the −1 oocyte and stopped when the oocyte completed its transition to zygote (Extended Data Fig. 1f). These results show that mitochondrial reduction is temporally coupled to OZT.

The specific timing of mitochondrial loss during OZT led us to hypothesize that mitochondrial reduction was a component of the OZT program triggered by sperm signaling. To test this, we blocked sperm signaling using RNAi knockdown of the sex determination gene *fem-3*, which inhibits sperm development in hermaphrodites^33^. Assessment of mitochondrial volume at the −1 oocyte relative to −2 oocyte revealed a complete abrogation of mitochondrial reduction in the −1 oocyte of feminized animals (Extended Data Fig. 1c,d). Measuring mitochondrial volume over time in the most proximal oocyte of feminized animals confirmed that there was no mitochondrial loss when OZT was blocked (Extended Data Fig. 1g).

We also observed that the absence of sperm altered mitochondrial morphology and distribution in the proximal oocytes of feminized animals. Rather than the tubular interconnected mitochondria observed in hermaphrodite oocytes, oocytes of feminized animals had small and donut-shaped mitochondria (Extended Data Fig. 1h). Oocyte development, maturation and fertilization is energetically demanding, and *C. elegans* oocyte mitochondria are organized as an interconnected branched network to support high ATP production^8^. Taken together, these results reveal a mitochondrial reduction event that is developmentally triggered by sperm signaling at the OZT that decreases mitochondrial transmission from mother to progeny. Our results also indicate that sperm induce networked, tubular mitochondrial morphology in oocytes prior to OZT.

### Macroautophagy activation during OZT

Given the substantial reduction of mitochondrial volume during OZT, we hypothesized that the oocyte could be using macroautophagy to eliminate mitochondria. Macroautophagy involves sequestration of cellular cargo by double membraned organelles called autophagosomes that then fuse with lysosomes to degrade cargo^34^.

To test if the autophagy-lysosome pathway was driving mitochondrial reduction at OZT, we disrupted key macroautophagy processes: autophagosome formation and maturation, autophagosome-lysosome fusion, and lysosomal acidification. We quantified mitochondrial reduction at OZT by measuring the ratio of mitochondrial signal in the zygote to that in the −2 oocyte, which is ∼0.5 in control conditions (∼2-fold reduction, Fig. 1c). RNAi knockdown was performed on genes responsible for autophagosome formation (*vps-34*, encodes a class III phosphoinositide 3-kinase that initiates autophagosome formation^34^) and autophagosome membrane elongation (*atg-16.1* and *atg-16.2*, parts of the ATG12-ATG5-ATG16L1 complex that promotes autophagosome vesicle expansion^34^). We also targeted VPS41 (Vacuolar protein sorting 41), a component of the homotypic fusion and protein sorting (HOPS) complex, which is required for autophagosome-lysosome fusion^35^ and disrupted lysosome function using RNAi knockdown of V-ATPase components^36^ (*vha-2*, *vha-9* and *vha-12*). Whereas there was a ∼2-fold reduction in mitochondrial signal between −2 oocyte and zygote in control conditions, knockdown of each of these genes abrogated the reduction in mitochondrial signal at OZT (Extended Data Fig. 2a,b).

We next asked whether macroautophagy activation is present and restricted to OZT by examining proteins marking autophagosome formation^37^ (endogenous ATG-9::GFP and mCherry::LC3) and lysosomes (endogenous CTNS-1::wrmScarlet) in oocytes. We found that ATG-9::GFP localizes as large puncta in the oocytes and that these ATG-9::GFP puncta were highly enriched at the −1 oocyte (over a ∼2-fold increase in number) compared to −2 oocytes (Extended Data Fig. 2c,f). We further assessed autophagosome number in oocytes, using germline expressed mCherry::LGG-2 (*C. elegans* ortholog of LC3, a key autophagosome membrane component), which also revealed upregulation in the −1 oocyte (Extended Data Fig. 2d,g). Lysosomes, measured using an endogenously tagged lysosomal membrane marker CTNS-1::wrmScarlet, were also increased in the −1 oocyte (Extended Data Fig. 2e,h).

Together, these results indicate that the oocyte activates the autophagy-lysosome system to degrade mitochondria during OZT. As macroautophagy of mitochondria is termed mitophagy^38^, we refer to this developmentally regulated degradation event as mitophagy at OZT (MOZT).

### MOZT requires mitochondrial fragmentation

Mitochondrial networks often undergo fragmentation to generate smaller, discrete mitochondria that are more conducive to autophagic degradation^39^. Therefore, we examined mitochondrial morphological changes at OZT by comparing mitochondrial network parameters (mitochondrial count, mean mitochondrial volume and sphericity) between oocytes and zygotes. We observed a mitochondrial fragmentation event during OZT—the zygote mitochondria were rounder, smaller and more discrete than oocyte mitochondria, which were tubular and interconnected (Fig. 2a,b). We conducted live imaging of the mitochondrial network during OZT and further confirmed dynamic mitochondrial fragmentation events (Fig. 2c and Supplementary Video 1, n=20/20 animals observed in the −1 oocyte). Average volume of the mitochondria was reduced from ∼2.0 µm³ (−2 oocyte) to ∼0.3 µm³ (zygote) during OZT (Fig. 2b), which was smaller than the −1 oocyte autophagosomes (∼0.6 µm³) (Extended Data Fig. 2i). These results demonstrate that oocyte mitochondria become fragmented during OZT.

**Fig.2:**
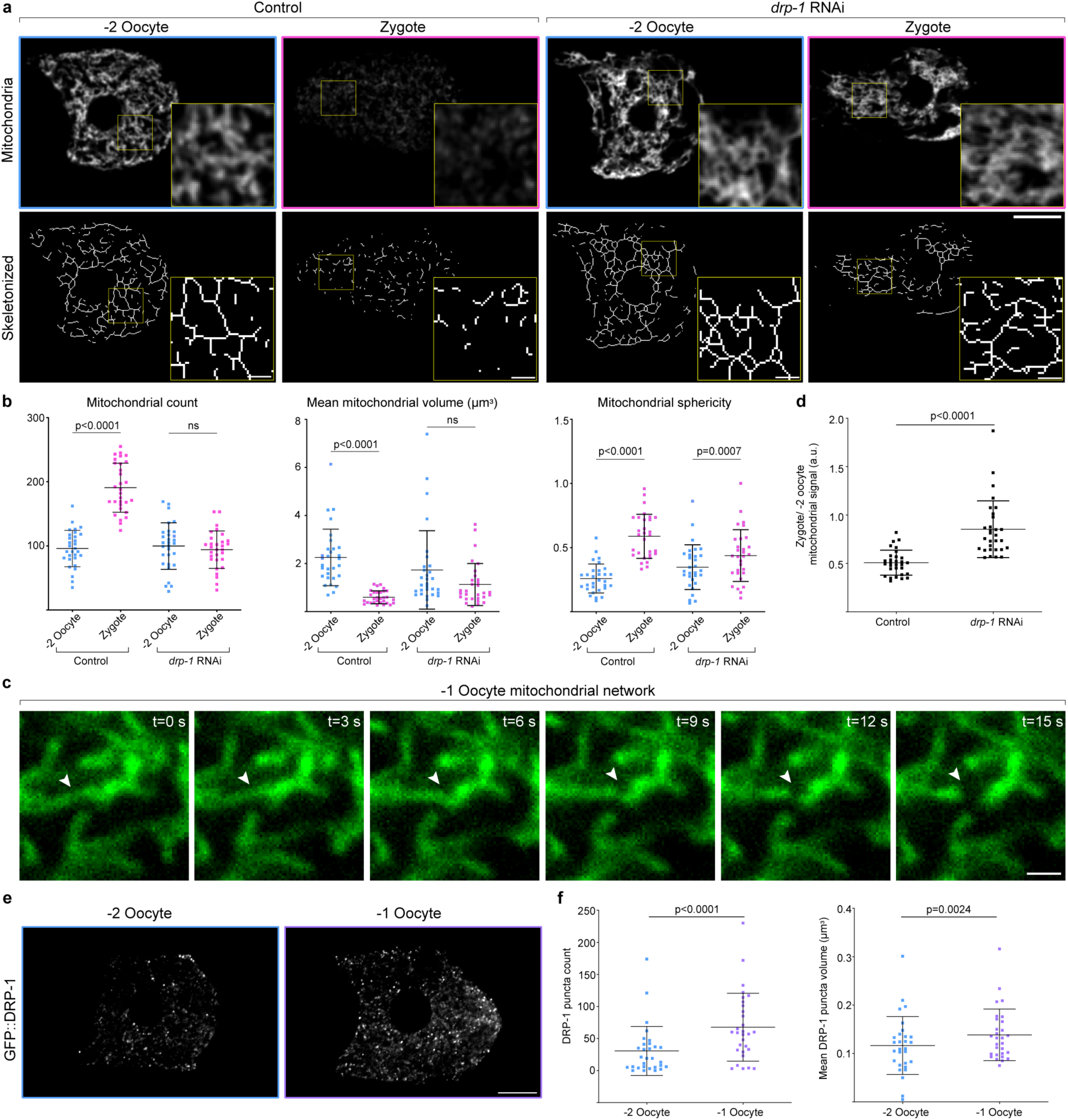
DRP-1 mediated mitochondrial fragmentation is necessary for MOZT. **a**, Top: Confocal fluorescence images showing the change in mitochondrial morphology (visualized with NDUV-2::mNG) between the −2 oocyte and zygote in an empty vector control (L4440) compared to a *drp-1* RNAi-treated animal. Scale bar, 10µm. Bottom: Skeletonized representation of mitochondria shown in upper panels. Boxed regions are magnified. Scale bars, 5 µm. **b**, Quantification of mitochondrial network parameters compared between −2 oocytes and zygotes in control and *drp-1* RNAi-treated animals. Data represent mean ± s.d. (n = 30 animals per condition) from three biological replicates. P values using two-tailed Student’s *t*-tests (mitochondrial count and sphericity) and Mann-Whitney test (mean mitochondrial volume). **c**, Time-lapse images of mitochondria (NDUV-2::mNG) undergoing fragmentation in the −1 oocyte. White arrows indicate the site of fragmentation on mitochondria. Example represents mitochondrial fragmentation events recorded at a single focal plane in the −1 oocyte and was observed in 20/20 animals over 4 min of observation in each. Scale bar, 2µm. **d**, Quantification of mitochondrial reduction during OZT in control vs *drp-1* RNAi-treated animals. Data represent ratio of total NDUV-2::mNG fluorescence intensity in zygote to that in the −2 oocyte for each animal. The data represent the mean ± s.d. (n = 30 animals) from three biological replicates. P value using a Mann-Whitney test. **e**, Endogenous GFP::DRP-1 puncta distribution in the −2 oocyte and −1 oocyte. Scale bar, 10µm. **f**, Increase in DRP-1 puncta number (left) and mean puncta volume (right) from −2 oocytes to −1 oocytes. Data represent mean ± s.d. (n = 30 animals) from three biological replicates. P values using Wilcoxon tests.

To uncover the molecular mechanism of mitochondrial fragmentation during OZT, we knocked down known effectors of mitochondrial morphology using RNAi and assayed for mitochondrial fragmentation at OZT in each condition (Supplementary Table 1). RNAi mediated knockdown of only one gene, *drp-1*, completely disrupted mitochondrial fragmentation during OZT, causing interconnected, elongated zygote mitochondria (Fig 2a,b). The *drp-1* gene encodes dynamin-related protein 1, a conserved large GTPase regulator of mitochondrial fission^40^. To test whether the mitochondrial fragmentation observed is essential for MOZT, we assessed for mitophagy after *drp-1* knockdown and found mitophagy was strongly inhibited (Fig. 2a,d). Next we used endogenously tagged GFP::DRP-1 to determine DRP-1 localization. GFP::DRP-1 was localized as puncta throughout the oocytes and there was a dramatic ∼2-fold increase in GFP::DRP-1 puncta number and ∼1.5-fold increase in puncta size in the −1 oocyte relative to the −2 oocyte (Fig. 2e,f). These findings indicate that oocytes upregulate DRP-1 specifically during the OZT to promote mitochondrial fragmentation for successful mitophagy.

Sustained mitochondrial fragmentation can initiate mitophagy in some cells^41^. Thus, we tested whether mitochondrial fragmentation is sufficient to trigger mitophagy in oocytes. We induced sustained mitochondrial fragmentation in oocytes using RNAi knockdown of *fzo-1*, the *C. elegans* ortholog of the mitochondrial fusion protein Mitofusin. Upon *fzo-1* knockdown the −2 oocyte mitochondria were fragmented, small and round compared to control oocytes (Extended Data Fig. 3a-c). However, the mitochondria in the −2 oocyte did not undergo mitophagy. Rather, mitophagy was still restricted to OZT (Extended Data Fig. 3d). These results indicate that mitochondrial fragmentation, while necessary for MOZT, is not sufficient to trigger mitophagy.

### Mitophagy receptor FUNDC1 mediates MOZT

Mitophagy is a selective form of autophagy that requires the assembly of mitophagy receptors on the outer mitochondrial membrane (OMM). After assembly, these receptors activate pathways to promote targeted mitochondrial degradation. The most studied are PINK1/Parkin-mediated and BNIP3-mediated mitophagy^38^, but a third, FUNDC1-mediated mitophagy, has been examined in response to hypoxia^30^.

Hence, we investigated whether the mitophagy receptors PINK-1 (*C. elegans pink-1*), BNIP3 (*dct-1*) or FUNDC1 (*fndc-1*) are necessary for MOZT. Examination of mitochondrial signal and volume as a measure of mitophagy at OZT in genetic null mutants of *pink-1* (*tm1779*) and *dct-1* (*luc194*) revealed no reduction in MOZT. In contrast, loss of *fndc-1* (*rny14*) showed complete inhibition of MOZT (Fig. 3a-c). In addition to promoting mitophagy, PINK1, BNIP3 and FUNDC1 regulate mitochondrial fragmentation^38,42,43^. Comparison of mitochondrial morphology at OZT in wild-type vs *fndc-1* animals showed that mitochondrial fragmentation during OZT was also significantly inhibited in the absence of FUNDC1 (Fig. 3d,e). These findings reveal a crucial role for FUNDC1 in mediating mitochondrial fragmentation and MOZT.

**Fig.3:**
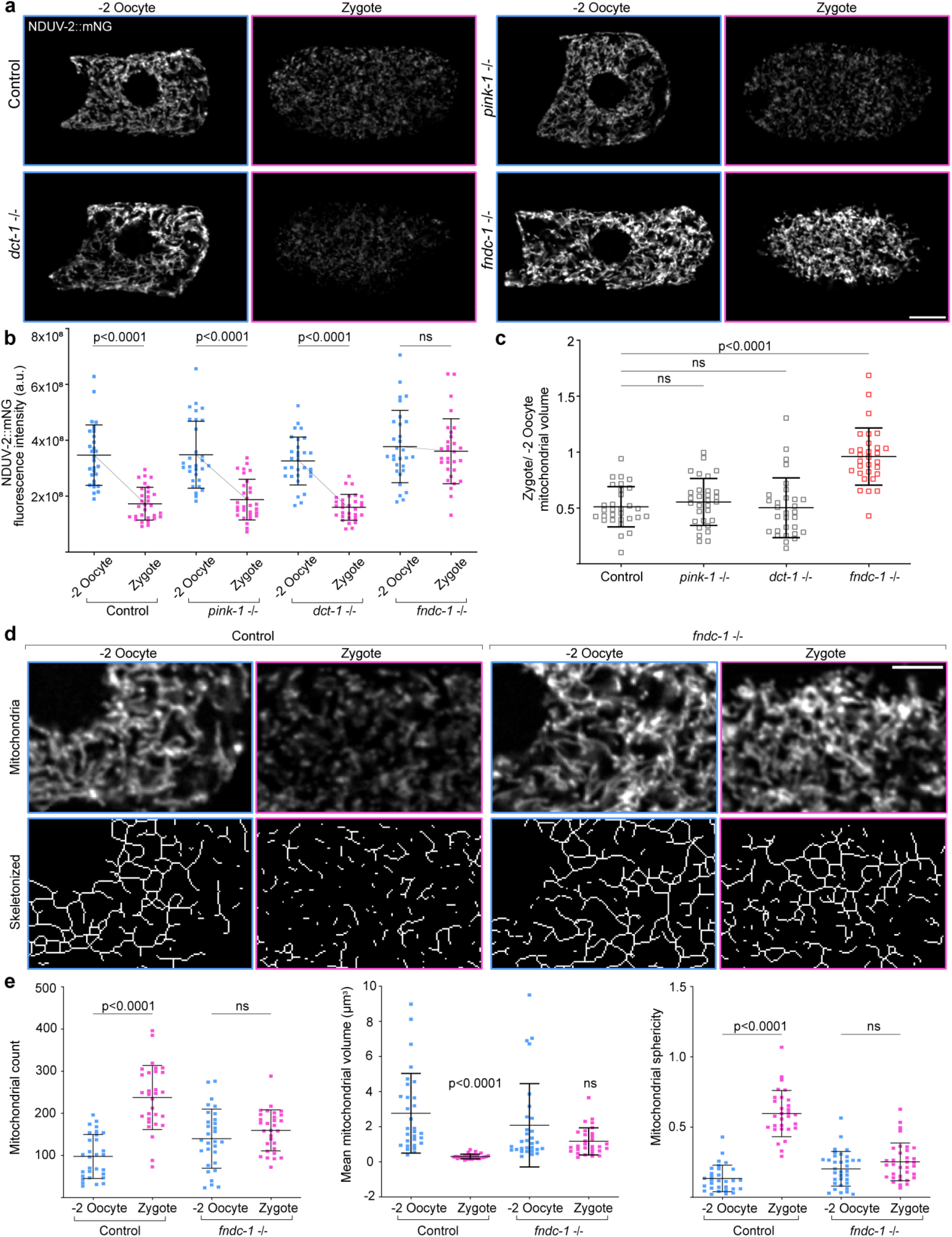
FUNDC1 is required for mitophagy and mitochondrial fragmentation during OZT. **a**, Confocal fluorescence images of mitochondria (visualized with NDUV-2::mNG) in the −2 oocyte and zygote of a control animal (wild-type) compared to null mutants of *pink-1*(*tm1779*), *dct-1*(*luc194*) and *fndc-1*(*rny14*). Scale bar, 10µm. **b**, Total NDUV-2::mNG fluorescence intensity compared between −2 oocytes and zygotes in control animals and null mutants of *pink-1*, *dct-1* and *fndc-1*. Data represent mean ± s.d. (n = 30 animals per strain) from three biological replicates. P values using two-tailed Student’s *t*-tests (control, *dct-1* and *fndc-1*) and Mann-Whitney test (*pink-1*). **c**, Quantification of reduction in mitochondrial volume during OZT in wild-type controls, *pink-1*, *dct-1* and *fndc-1* mutant animals. Data represent ratio of mitochondrial volume in zygote to that in −2 oocyte for each animal. Data represent mean ± s.d. (n = 30 animals per strain) from three biological replicates. P values using a Kruskal Wallis followed by Dunn’s multiple comparisons test. **d**, Top: Confocal fluorescence images showing the change in mitochondrial morphology (NDUV-2::mNG) between the −2 oocyte and zygote in a wild-type control compared to a *fndc-1* mutant animal. Bottom: Skeletonized representation of mitochondria shown in upper panels. Scale bars, 5µm. **e**, Quantification of mitochondrial network parameters compared between −2 oocytes and zygotes in a wild-type control and *fndc-1* mutant animals. Data represent mean ± s.d. (n = 30 animals per strain) from three biological replicates. P values using two-tailed Student’s *t*-tests and Mann-Whitney tests, depending on normality of distribution. Supplementary Fig.1 shows uncropped fluorescence images.

### FUNDC1 marks mitochondrial fission sites

FUNDC1, a conserved outer mitochondrial membrane protein, both mediates selective autophagy and accumulates at mitochondrial associated membranes (MAMs) in response to hypoxia, to recruit DRP-1 for mitochondrial fragmentation^43^. We next sought to determine FUNDC1 localization during OZT using an endogenous mRb3::FNDC-1 knock-in, in combination with the endogenous mitochondrial protein NDUV-2::mNG. In −2 and −1 oocytes, mRb3::FNDC-1 signal was localized diffusely to the mitochondrial membrane (Fig. 4a), consistent with OMM localization. We also observed variable large puncta (Fig. 4a), which increased in number and signal intensity in −1 oocytes (Fig. 4b,c). As FUNDC1 promotes selective mitophagy through its LIR (LC3 interacting region) that facilitates mitochondrial targeting to autophagosomes, and also promotes mitochondrial fragmentation by recruiting DRP-1 through a cytosolic loop^43^, we next wanted to examine the relationship between FUNDC1 punctae and mitochondrial dynamics.

**Fig.4:**
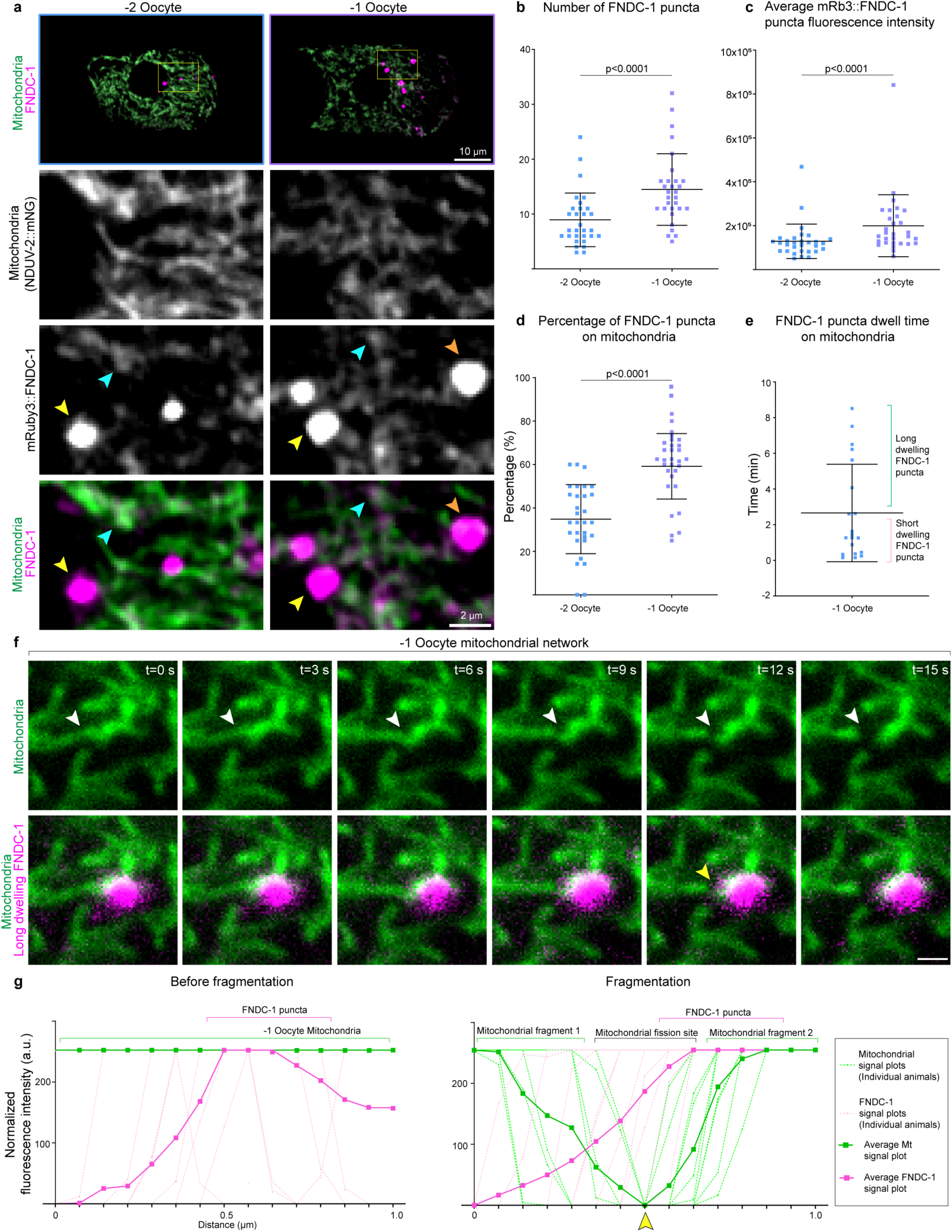
Stable FUNDC1 punctae localize at sites of mitochondrial fragmentation during OZT. **a**, Top: Merged confocal fluorescence images of mitochondria (NDUV-2::mNG) and mRb3::FNDC-1 in the −2 oocyte and −1 oocyte. Scale bar, 10µm. Bottom: Magnifications of boxed regions. Blue arrows indicate mRb3::FNDC-1 signal localized at the outer mitochondrial membrane (OMM). Orange arrows indicate mRb3::FNDC-1 puncta associated with the mitochondrial network. Yellow arrows indicate mRb3::FNDC-1 punctae adjacent to the mitochondrial network. Scale bar, 2µm. **b**,**c**, Number (**b**) and mean fluorescence intensity (**c**) of mRb3::FNDC-1 puncta in −2 oocytes and −1 oocytes. Data represent mean ± s.d. (n = 30 animals per strain) from three biological replicates. P value using a Wilcoxon test. **d**, Proportion of mRb3::FNDC-1 puncta localized on mitochondria in −2 oocytes and −1 oocytes. Data represent mean ± s.d. (n = 30 animals per strain) from three biological replicates. P value using a two-tailed paired Student’s *t*-test. **e**, Duration for which mRb3::FNDC-1 puncta localized on mitochondria in −1 oocytes. Data represent mean ± s.d. (n = 20 mRb3::FNDC-1 puncta from 12 animals) from four independent experiments. Distribution of dwell time data represents two distinct behaviors of mRb3::FNDC-1 puncta with respect to mitochondrial association. **f**, Top: Time-lapse images of mitochondria (NDUV-2::mNG) undergoing fragmentation in the −1 oocyte. White arrows indicate site of fragmentation on mitochondria. Bottom: Merged fluorescence images showing mRb3::FNDC-1 puncta localize at the mitochondrial fragmentation site. Scale bar, 2µm. Data is representative of n = 10/10 FUNDC1 puncta stabilized on −1 oocyte mitochondria for more than 4 min. Supplementary Video 1 shows the time-lapse data from which **f** is derived. **g**, Graphs are line scans (1µm) drawn across FUNDC1 puncta stabilized on mitochondrial surface in the −1 oocyte and show the normalized fluorescence intensity of mitochondria (green) and FUNDC1 puncta (magenta). Left: Line scans along mitochondrial regions that are associated with FUNDC1 puncta before fragmentation. Right: Line scans along same mitochondrial regions, showing break in mitochondria occurs at FUNDC1 puncta co-localized for t > 4 min. Yellow arrow at fragmentation site on representative image from **f** corresponds to yellow arrow shown on the line scan. Data represents line scans from 10 independent experiments shown as well as the average line scans.

MAMs are formed at the interface between ER and mitochondria^43^ and biochemical analyses have shown that FUNDC1 accumulates at MAMs^43^. Examination of mRb3::FNDC-1 puncta in tandem with an ER marker (GFP::SP12) and mitochondrial marker (NDUV-2::mNG) revealed the mRb3::FNDC-1 puncta were always on or adjacent to the ER and mitochondria, consistent with MAM localization (n=20/20, Extended Data Fig. 4a,b). Proper balance of mitochondrial fission-fusion is required for MAM formation^44^. We disrupted the formation of MAMs by inducing hyperfusion of mitochondria (*drp-1* RNAi) and noted that in the −1 oocytes, mRb3::FNDC-1 puncta were now sparse in number compared to control (Extended Data Fig. 4c,d). Moreover, mRb3::FNDC-1 signal enrichment shifted from punctate localization to OMM (non-punctate) localization (Extended Data Fig. 4e,f). We also noted that in −2 oocytes, ∼35% of mRb3::FNDC-1 puncta were clearly co-localized at specific foci on the surface of mitochondria and this proportion increased to ∼60% in the −1 oocytes (Fig. 4d). Live-cell imaging of mRb3::FNDC-1 puncta together with mitochondria (NDUV-2::mNG) in −1 oocytes showed that dwell time of the puncta on mitochondria was diverse, ranging from 10 s to 8.6 min (Fig. 4e).

Tracking of FUNDC1 puncta revealed two distinct behaviors (Fig. 4e, Extended Data Fig. 5a and Supplementary Video 2): ∼70% of observed puncta (n=20 puncta from 12 animals) co-localized on mitochondria for short time frames ranging from 10 s to 2 min (Fig. 4e) and there was no noticeable effect on the observed mitochondrion (Extended Data Fig. 5a,b,d, n = 20/20). In contrast, ∼30% of FUNDC1 puncta (n=20) were localized on mitochondria for longer timeframes ranging from 4 min to 8.6 min (Fig. 4e) and this always led to a mitochondrial fragmentation event (Supplementary Video 1, Fig. 4e,f,g, n=10/10 and Extended Data Fig. 5c,d). These results suggest that stable FUNDC1 aggregates on oocyte mitochondria promote mitochondrial fragmentation, which is required for MOZT (Extended Data Fig. 4g).

### FUNDC1 responds to mitochondrial stress

Previous studies in cancer cells, mice cardiomyocytes and *C. elegans* body wall muscle have shown that cells respond to hypoxic stress by accumulating FUNDC1 at MAMs^30,45^. Whether cells utilize FUNDC1 pathway in response to non-hypoxic stimuli that induce mitochondrial damage is unknown.

To test this, we first induced lesions in the *C. elegans* oocyte mtDNA using UVC treatment, which creates cyclobutane pyrimidine dimers in mtDNA that can lead to mtDNA mutations during replication^46^. UVC-induced mtDNA damage in oocytes resulted in a dramatic upregulation of mRb3::FNDC-1 signal on mitochondrial membranes, increased the number of MAM-associated punctae, and elevated accumulation of mRb3::FNDC-1 signal at the MAMs/punctae (Fig. 5a-d). Besides UVC-induced mtDNA damage, we used a *C. elegans* strain harboring mtDNA with a large deletion (3054-bp deletion) called the *uaDf5* deletion, which models large mtDNA deletions that are commonly detected in animal oocytes, including humans^6,25–27,47^ (Extended Data Fig. 6a). We found that in the presence of the *uaDf5* deletion, oocytes also exhibited elevated levels of mRb3::FNDC-1 both on the mitochondrial membranes and in puncta associated with MAMs (Extended Data Fig. 6b-e).

**Fig.5:**
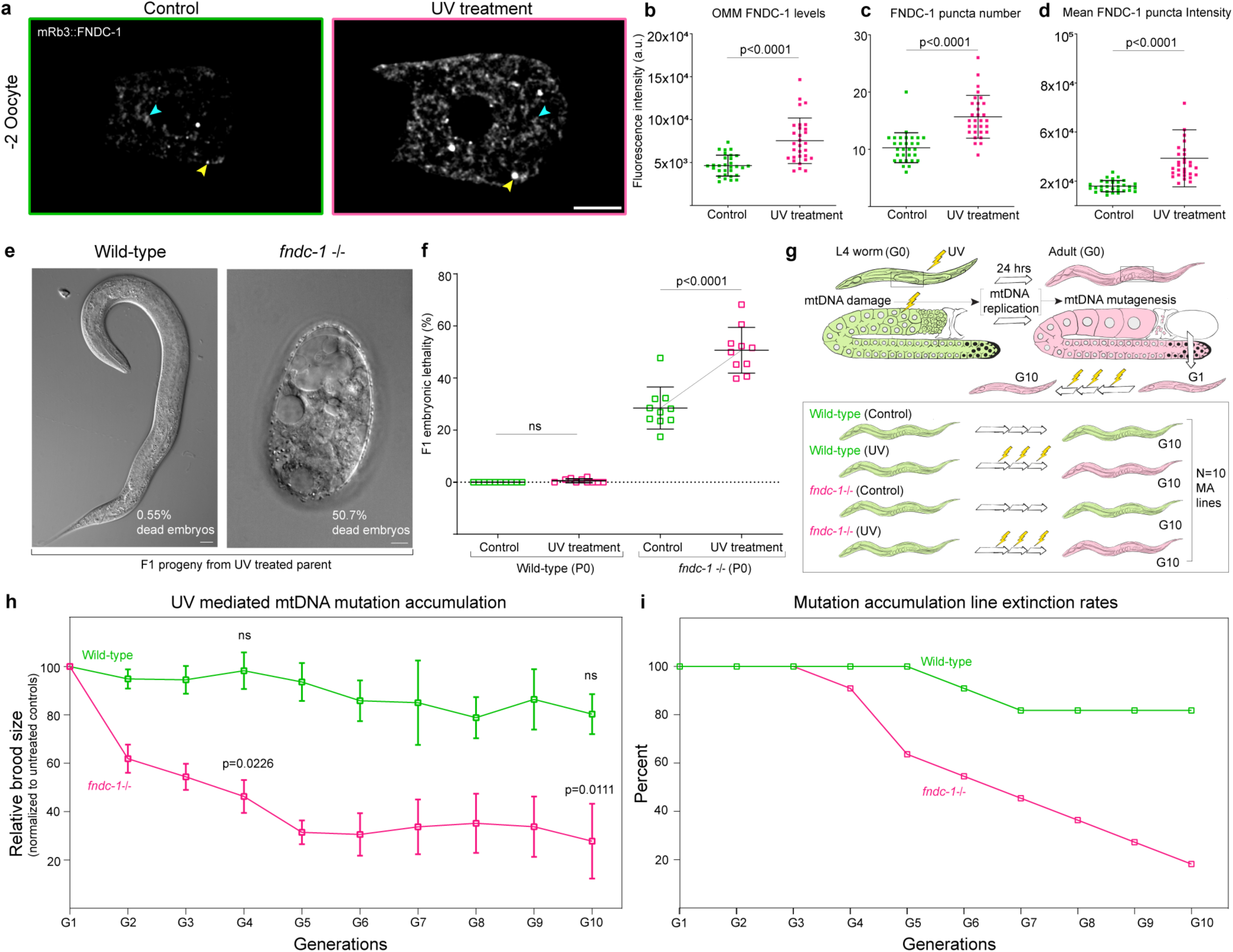
Loss of FUNDC1 leads to increased embryonic lethality and population extinction upon mtDNA damage accumulation. **a**, Endogenous mRb3::FNDC-1 distribution in the −2 oocyte of a control animal compared to a UVC treated animal. Blue and yellow arrows indicate OMM and mitochondria associated membrane (MAM) localized mRb3::FNDC-1 signal, respectively. Scale bar, 10µm. **b**,**c**,**d**, Mean fluorescence intensity of OMM localized mRb3::FNDC-1 (**b**); number (**c**) and mean fluorescence intensity (**d**) of mRb3::FNDC-1 puncta in −2 oocytes of control animals compared to UVC treated animals. Data represent mean ± s.d. (n = 30 animals per condition) from three biological replicates. P values using two-tailed Student’s *t*-test (**b**) and Mann-Whitney tests (**c**,**d**). **e**, DIC images of representative F1 progeny of wild-type and *fndc-1* mutant parents (P0) exposed to UVC treatment (exposure at L4 stage). Scale bars, 10µm. Mean embryonic lethality percentage ± s.d. among F1 progeny is indicated. **f**, Embryonic lethality (mean ± s.d.) in F1 progeny of wild-type and *fndc-1* mutant parents in control conditions compared to UVC treatment conditions. n = 10 animals were used as P0 parents per strain per condition in three independent experiments. P values using a Wilcoxon test (wild-type) and a two-tailed Student’s *t*-test (*fndc-1*). **g**, Schematic of mutation accumulation (MA) experimental design. Green animals denote undamaged worms, and magenta animals denote worms with mtDNA damage. **h**, Relative brood size as indicator of reproductive fitness of MA lines across generations. Total brood size was calculated at each generation (G0 to G10) in UV MA lines and normalized against that of respective control MA lines. Lines (green, wild-type UV MA lines; magenta, *fndc-1* UV MA lines) indicate decline in reproductive fitness relative to controls across generational time upon UVC treatment. Data points represent mean ± s.e.m (n = 10 animals per strain per treatment) from three independent experiments. P values using Kruskal-Wallis test followed by Dunn’s multiple comparisons test. **i**, Extinction rates of wild-type and *fndc-1* UV MA lines. Y-axis denotes the percentage of MA lines present at each generation out of 10 wild-type UV MA lines (green line) and 10 *fndc-1* UV MA lines (magenta line).

Lastly, we subjected animals to various exogenous mitochondrial damaging agents-oxidative damage through paraquat treatment, mitochondrial uncoupling through FCCP treatment, and the recently identified mitochondrial toxicant 6PPD-Q (*N*-(1,3-dimethylbutyl)-*N*′-phenyl-*p*-phenylenediamine-quinone, rubber tire oxidation product), which causes mitochondrial dysfunction in *C. elegans* by affecting mitochondrial complexes I and II^48^. Upon FCCP and 6PPD-Q treatment, we observed increased FUNDC1 expression in oocytes (Extended Data Fig. 7a-c). Together, this indicates that oocytes upregulate FUNDC1 in response to diverse types of mitochondrial insult.

Given the role of FUNDC1 in mitochondrial fragmentation and mitophagy and the substantial increase in FUNDC1 expression in the oocyte upon mitochondrial damage, we asked if mitochondrial damage led to increased mitochondrial fragmentation and mitophagy at OZT. We found that mitochondrial fragmentation and mitophagy, however, were unchanged even upon the introduction of deleterious mtDNA through UVC treatment (Extended Data Fig. 7d-f). We also found that the number of ATG-9::GFP vesicles, an indicator of autophagosome biogenesis^37^ was not elevated upon UVC treatment (Extended Data Fig. 7g-j). These findings indicate that the amount of fragmentation and mitophagy at OZT is not determined by level of mitochondrial damage. Instead, our results suggest oocyte mitophagy levels might be set at a specific developmentally regulated level and that increased FUNDC1 expression upon mitochondrial damage could be functioning to promote selectivity against damaged mitochondria during MOZT (*i.e.*, selecting which mitochondria are targeted for degradation).

### FUNDC1 mediates cross-generational purifying selection

To test if FUNDC1-mediated MOZT could be a cross-generational mitochondrial quality control event that removes damaged mitochondria, we examined transmission of the *C. elegans* mtDNA deletion *uaDf5*. The *uaDf5* deletion serves as a dynamic indicator of mtDNA quality control. When mtDNA is replicated, the mtDNA copies harboring the *uaDf5* deletion has a selfish replication advantage and increases in prevalence over wild-type mtDNA. However, the deletion never completely dominates because mitochondrial quality control mechanisms counteract the *uaDf5* deletion^49,50^. If FUNDC1 acts against deleterious mtDNA, we would expect the levels of *uaDf5* to be higher in progeny of *fndc-1* mutant mothers compared to wild-type progeny.

Interestingly, previous studies have shown a ∼13% decrease in *uaDf5* heteroplasmy levels between wild-type mothers and L1 larval offspring, which is consistent with a mtDNA quality control process during the mother-to-offspring transition^23^. We first examined *uaDf5* heteroplasmy levels in adult parent germlines in wild-type and *fndc-1* mutant backgrounds and found that the levels of *uaDf5* heteroplasmy were the same (Extended Data Fig. 6f). This indicates that mtDNA heteroplasmy in the adult germline is not under regulation by FUNDC1. We then examined the levels of *uaDf5* heteroplasmy in the L1 larval progeny and found that *uaDf5* heteroplasmy was ∼13% higher in *fndc-1* L1 larval progeny compared to wild-type progeny (Extended Data Fig. 6g). Together, these results reveal that FUNDC1 is required to reduce the load of deleterious mtDNA inherited by *C. elegans* progeny, indicating FUNDC1-mediated MOZT could be a key purifying selection event that resets mitochondrial health during mother-to-offspring transmission.

### FUNDC1 promotes species endurance

The increased amounts of deleterious mtDNA mutations inherited in *fndc-1* mutant animals, could have severe detrimental effects when accumulated over several generations^13,51^. To test this, we used UVC exposure to generate mtDNA damage in animals without FUNDC1. In untreated *fndc-1* null animals, we noted ∼25% embryonic lethality (Fig. 5e,f). After UVC induced oocyte mitochondrial damage, however, we observed ∼50% embryonic lethality in progeny (F1), whereas in wild-type animals there was no increase (Fig. 5e,f).

We next investigated the detrimental effects of repeated mtDNA damage over multiple generations in wild-type and *fndc-1* animals through a mutation accumulation (MA) experiment^52^. UVC treated MA lines and untreated MA lines were generated in wild-type and *fndc-1* null mutants. In the UVC mtDNA mutagenesis condition, for every generation a single L4 animal was selected as a progenitor for the next generation and was treated with a 25J/m² dose of UVC radiation to induce mtDNA mutations in the oocyte mitochondria for 10 sublines for 10 generations (methods, Fig. 5g). Reproductive fitness declined at a rapid rate in the *fndc-1* UV MA lines compared to wild-type UV MA lines (Fig. 5h). Upon just one generation of UVC treatment (G2), *fndc-1* animals exhibited a dramatic ∼40% decline in fecundity, while wild-type control animals had no decrease (Fig. 5h). After 10 generations of mutation accumulation, wild-type UV MA lines exhibited an insignificant decline in reproductive fitness compared to untreated wild-type MA lines, while *fndc-1* UV MA lines showed severely reduced fecundity (∼75% decline) (Fig. 5h). Moreover, by the end of 10 generations of MA, only 19% of wild-type UV MA lines were extinct, while 82% of *fndc-1* UV MA lines were extinct (Fig. 5i).

To rule out deteriorating sperm quality over the 10 generations causing population decline, we crossed *fndc-1* UV MA lines (fourth generation, G4) with untreated wild-type males and assessed if the introduction of healthy sperm improved fecundity (see methods). However, there was no significant improvement to reproductive fitness of the *fndc-1* UV MA lines (Extended Data Fig. 8a-c), indicating that the decreased fecundity in *fndc-1* UV MA lines is not a paternal effect.

To test whether the protective quality control of FUNDC1 was acting during MOZT, we expressed mRb3::FNDC-1 specifically after fertilization in the embryos derived from *fndc-1* UV MA line mothers (G4, see Methods) and found that embryonic expression of FUNDC1 did not reduce embryonic lethality (Extended Data Fig. 8d-f), offering compelling evidence that FNDC-1/FUNDC1 acts in the oocyte, consistent with a role at MOZT.

Finally, we examined mitochondrial respiration parameters in G4 MA lines to determine if there were any detrimental effects from UVC induced mtDNA mutation accumulation on mitochondrial function (Extended Data Fig. 9). Adults from the *fndc-1* UV MA lines exhibited significantly increased non-mitochondrial OCR compared to wild-type UV MA lines (Extended Data Fig. 9), which can occur due to increased enzymatic production of reactive oxygen species (ROS), a negative indicator of bioenergetic health^53^. We conclude that maternal FUNDC1 acting in the late oocyte is important for its function in maintaining mitochondrial and reproductive fitness across multiple generations, thus contributing to species endurance (Extended Data Fig. 10).

## Discussion

Mature oocytes are metabolically active and thus contain the largest concentration of mitochondria of any cell in the organism^22^. Mitochondrial activity is maintained at high levels in the oocyte for energy production and synthesis of biomolecules essential for fertilization and embryogenesis^54^. However, mitochondrial oxidative metabolism generates ROS as by-products, that at high concentrations cause mtDNA damage and mitochondrial dysfunction^55^. In addition, oocytes accumulate protein aggregates and oxidated proteins, furthering mitochondrial damage^56–58^. Oxidative mtDNA lesions, large mtDNA deletions and rearrangements are commonly detected in mature oocytes, indicative of mitochondrial damage^25–28^. When present at high levels, these diminish fertilization potential of the oocyte and causes embryonic developmental defects^59^. The mechanisms by which oocytes balance maintaining mitochondrial abundance and activity with transmission of healthy mitochondria to the zygote and the next generation are unknown. In this study we discovered MOZT (mitophagy at OZT), a developmentally programmed FUNDC1 receptor-mediated mitochondrial quality control event that allows oocytes to maintain this balance.

Developmentally programmed mitophagy events occur in cells like keratinocytes and erythrocytes and are mediated by the mitophagy receptors NIX and BNIP3^60^. MOZT is unique from known programmed mitophagies in its regulation by FUNDC1. While NIX and BNIP3 have BH3 domains allowing them to interact with apoptosis regulators like Bcl-2^60^, FUNDC1 contains domains that bind to mitochondrial fission-fusion proteins like DRP-1^43^. *C. elegans* oocytes have elongated interconnected mitochondria to support high energy production^8^. We found that as oocytes undergo OZT, they increasingly accumulate FUNDC1 at MAMs and upregulate DRP-1 for rapid mitochondrial fragmentation. Through live imaging, we showed that FUNDC1 is highly dynamic, but concentrations of FUNDC1 at MAMs stably localize on mitochondrial surfaces and promote mitochondrial fragmentation in a DRP-1 dependent manner during OZT. Our findings support reports showing that overexpression of FUNDC1 in *C. elegans* body wall muscle increases FUNDC1 puncta and mitochondrial fragmentation^45^. We found that loss of DRP-1 inhibits MOZT, strongly supporting the notion that mitochondrial fragmentation is a prerequisite for programmed mitophagy. Furthermore, our work suggests that FUNDC1 could be the primary mitophagy regulator in cells with highly interconnected mitochondrial networks^61^, due to their MAM localization and close association with mitochondrial fission proteins.

Maternally inherited mtDNA mutations in mice lead to fertility decline^62^ and accumulation of deleterious mtDNA in *C. elegans* upon mtDNA 6mA hypomethylation has been reported to lead to sterility in 12 generations^63^. Thus, mtDNA quality control during mother-to-offspring transmission is crucial for species survival. Multiple mechanisms regulate mtDNA quality during germline and oocyte development. PINK1 functions independent of mitophagy and promotes mtDNA quality control in the primordial germ cells in *C. elegans*^64^ and during oogenesis in *Drosophila*^3,65^ by selectively preventing the replication of defective mtDNA. In addition, programmed mitophagy occurs during early oogenesis in *Drosophila*, regulated by the mitophagy receptor BNIP3^5^. The programmed cell death pathway also mediates mtDNA quality control during *C. elegans* germline aging^23^. Yet, none of these mechanisms occur during the long period of oocyte growth and differentiation when metabolic activity is highest and thus do not ensure the transmission of healthy mitochondria to offspring^29^. Our findings support FUNDC1-mediated MOZT as the counteracting force against maternal transmission of deleterious mtDNA that functions at the last step in oocyte development. We found that FUNDC1 was specifically upregulated during MOZT, and our results indicate it promotes mtDNA quality control during this time. Using *uaDf5*, a large deletion in mtDNA that exists with wild-type mtDNA and thus allows detection of mtDNA quality control, we found that adult germlines of *fndc-1* null mutant parents had similar levels of *uaDf5* compared to wild-type parent germlines. However, the L1 larval progeny of *fndc-1* null mutants inherited ∼13% increased abundance of mtDNA deletion (*uaDf5*), with some larvae having unprecedented proportions of the *uaDf5* deletion (70-90% of mitochondrial genome). These findings strongly suggest that FUNDC1 mitophagy acts at the maternal-to-offspring transition to ensure mtDNA quality control across generations.

We also found that MOZT preserves female fertility and progeny health against mtDNA damage. UVC mediated mtDNA damage in maternal germlines of *fndc-1* null mutants resulted in a ∼2-fold increased embryonic lethality in the progeny. Further supporting the time of action of FUNDC1 at OZT, embryonic expression of FUNDC1 did not rescue this lethality. Upon several generations of mtDNA mutation accumulation, *fndc-1* null animals experienced severe declines in fertility. Four generations of UV induced mtDNA mutation accumulation reduced fecundity of *fndc-1* null animals by ∼75% and led to impaired bioenergetics in the whole animal, indicated by increased non-mitochondrial oxygen consumption^53^. With just 10 generations of mutation accumulation (MA), we observed 82% extinction of animal populations lacking FUNDC1.

Neither PINK1, BNIP3 mitophagy nor programmed cell death has a role in preventing accumulation of deleterious mtDNA across generations^52,23^ indicating FUNDC1-mediated MOZT could be the primary intergenerational mtDNA quality control mechanism that promotes species continuity.

FUNDC1 levels are significantly upregulated specifically during OZT to promote mitochondrial fragmentation and MOZT, indicating a developmental program. Yet, we also found that oocytes upregulate endogenous FUNDC1 expression further, after exposure to a broad range of mitochondrial insults, including mtDNA lesions, mtDNA deletions, uncoupling damage through FCCP and mitochondrial dysfunction. This suggests FUNDC1 is also adaptive to environmental insults and is a component of a previously unknown oocyte mitochondrial damage sensing mechanism. Interestingly, oocytes do not further activate mitophagy or mitochondrial fragmentation upon presence of deleterious mtDNA, but rather undergoes MOZT to the same extent as untreated conditions. Increased levels of FUNDC1 on mitochondrial surface typically promotes autophagosome recruitment to the mitochondrion through a classical LIR motif (LC3 interacting region) on FUNDC1^43^. Thus, the increased FUNDC1 expression at OMM in oocytes upon mitochondrial damage might serve as the basis for selectivity during autophagic degradation at MOZT. Our findings that oocytes upregulate FUNDC1 levels in presence of *uaDf5* support this notion, since we show that FUNDC1 mitophagy selects against *uaDf5*. As FUNDC1 also promotes mitochondrial and cellular Ca^2+^ homeostasis by regulating the ER-mitochondrial Ca^2+^ exchanger IP3R3 at MAMs^66^, upregulation could also mediate mitochondrial and cellular homeostasis under mitochondrial damage conditions.

The consistency in the level of mitochondrial reduction (∼2-fold) even upon mitochondrial insult reveals that a critical threshold of mitochondria is precisely set to be inherited by the zygote at the end of MOZT. We found that macroautophagy is not upregulated further upon mtDNA mutagenesis, indicating that the threshold for mitochondrial degradation could be limited by the autophagy-lysosome system. As embryogenesis requires a critical level of mtDNA be inherited from the mature oocyte^67,68^, a threshold might have evolved that balances mtDNA quality control and metabolic needs of the embryo. During OZT, oocytes also transition from relying on mitochondrial oxidative phosphorylation to glycogen metabolism. Early embryo mitochondria are relatively dormant in function, displaying reduced membrane potential, indicating organelle and metabolic rewiring^69^. Reduction of mitochondria through MOZT could also help promote the shift in cellular metabolism at OZT. Consistent with this idea, programmed mitophagy promotes precise remodeling of mitochondrial proteome and cellular metabolism during neurogenesis and heart development and is crucial for a metabolic switch to glycolysis during retinal ganglion cell differentiation^70^.

mtDNA copy numbers drop during OZT in zebrafish^71^, mice^72^, cow^22^ and humans^73^. Furthermore, mtDNA mutations and deletions occur at significant levels in late oocytes in many different mammalian species, which are removed upon fertilization. These observations have suggested the existence of a cryptic purifying selection mechanism^6,7,28^. Intriguingly, mature oocytes from fish, amphibian and mammalian species, including humans, express FUNDC1^74–78^. Together, this raises the possibility that MOZT could be a conserved developmental program crucial in preserving mitochondrial health during maternal inheritance.

## Acknowledgements

We thank the members of the laboratory of D.S. and J.M. for helpful discussions, Adam Soh and Lena Basta for reading and editing of the manuscript, and Litong Yang for help with maintaining worm strains. We thank Amy Wehman for the WEH722 strain and Patrick Laurent for the PHX5270 strain. We also thank Laura Jameson for technical help with mitochondrial toxicants. This work was supported by R35GM118049 to D.R.S, P42ES010356 and T32ES021432 to J.M.

## Author contributions

S.B.T. conceived the project, designed the study, performed most of the experiments, analyzed the data, wrote the manuscript and prepared the figures. S.B. performed the qPCRs for measurement of *uaDf5* ratios and helped optimize dosages for UVC treatment under supervision of J.M. K.M. performed measurements of mitochondrial respiration in MA lines and analyzed the data under supervision of J.M. Q.C. and I.K. constructed molecular biology tools for the project. D.S. supervised the project, edited the manuscript and acquired funding for the project.

## Supplementary Information

Supplementary information is available for this paper. Refer to Supplementary Tables 1-3, Supplementary Fig. 1-3 and Supplementary Videos 1, 2.

## Corresponding authors

Correspondence and requests for materials should be addressed to David R. Sherwood.

## Methods

### *C. elegans* strains and culture conditions

*C. elegans* strain were fed *Escherichia coli* OP50 and raised under standard conditions at 20°C. N2 Bristol strain was used as the wild-type strain^79^. All oocyte-to-zygote transitions (OZT) observed and imaged were in day 1 adult hermaphrodites, 24 h after L4 stage. All *C. elegans* strains and alleles used in this study are listed in Supplementary Table 2.

### Strain generation through genome-editing

The fluorescent protein mNeonGreen was knocked-in at the C-terminus of endogenous coding sequence of mitochondrial proteins TOMM-20 and NDUV-2, using CRISPR-Cas9-mediated genome editing as previously described before^80,81^. Respective sgRNAs (sgRNA for TOMM-20: 5′-GACACCGACGACTTGGAGTAA-3′, sgRNA for NDUV-2: 5′-GGCTGCTCTTAAATAAACGCT-3′) and repair template plasmids with the *mNeonGreen (mNG)* coding sequence containing homology arms were used. After injection, recombinant animals were identified in the F3 offspring based on selectable markers (hygromycin B resistance and dominant-negative *sqt-1 rol* phenotype). In positive candidates, selection markers were excised from the genome through Cre-Lox recombination. The strains generated were outcrossed with N2 wild-type strain for four generations before further experimental use. Primers used for genotyping are included in Supplementary Table 3.

### Strain generation through genetic crosses

The following *C. elegans* strains were created using standard mating techniques: NK3285 (*qy174(nduv-2::mNG)V; rny15(mRuby3::fndc-1*^82^*)II*); NK3286 (*ojIs23[pie-1p::GFP::SP12, unc-119(+)]; rny15(mRuby3::fndc-1)II*); NK3282 (*qy174 (nduv-2::mNG)V; pink-1(tm1779*^83^*)II*); NK3283 (*qy174 (nduv-2::mNG)V; dct-1(luc194*^84^*)X*); NK3284 (*qy174 (nduv-2::mNG)V; fndc-1(rny14*^82^*)II*); NK3287 (*rny15(mRuby3::fndc-1)II; uaDf5/+* mtDNA); NK3288 (*fndc-1(rny14)II; uaDf5/+* mtDNA). To construct strains with *rny14* and *rny15* heteroplasmic for *uaDf5* mtDNA (*uaDf5/+*), 10-15 males homozygous for *rny14* and *rny15* were used to set up genetic cross with 3-5 hermaphrodites heteroplasmic for *uaDf5* mtDNA. Animals homozygous for *rny14* and *rny15* alleles were selected among the F3 progeny. Retention of *uaDf5* mtDNA in resulting crosses was confirmed by PCR. Oligos used in this study for genotyping analyses are listed in Supplementary Table 3.

### Cloning of *fndc-1* for embryonic expression

The embryonic expression of mRb3::FNDC-1 was achieved by construction of the transgene *pie-1p::mRuby3::fndc-1::unc-54* 3’UTR. The *pie-1* promoter (1.114-kb fragment), full genomic sequence of *fndc-1* (535-bp) and the *unc-34* 3’UTR (868-bp fragment) were fused in the pAP088 plasmid. The construct was injected into the gonads of at least 20 young adults, two from each of the 10 different *fndc-1* (UV) MA lines (G4) along with 2.5ng/mL co-injection markers (*myo-2p::gfp* and *myo-3::gfp*). The construct also contained coding sequence of a dominant negative *sqt-1* mutation that gives rise to a *rol* phenotype when expressed. Viability assays and fluorescence imaging were performed on F1 offsprings from successful injections, identified by presence of mRb3::FNDC-1 and selectable markers (green pharynx, green body wall muscle and dominant negative *sqt-1 rol* phenotype).

### Microscopy and image analysis

Differential interference contrast images and confocal fluorescence images were acquired on an upright Carl Zeiss Axio Imager A1 microscope attached to a Yokogawa CSU-W1 spinning disk confocal head. Imaging experiments were performed using a Zeiss 63x Plan-Apochromat (NA 1.40) oil immersion objective and a Hamamatsu ORCA-Fusion or a Hamamatsu ORCA-Quest camera with 488nm, 505nm and 561nm lasers. For imaging OZT, day 1 adult worms were anaesthetized in a 2μL solution of 5mM Levamisole placed onto the center of 5% agar pads and a #1.5 cover slip was placed on top before imaging at room temperature. For time-lapsing experiments, the slide was sealed using VALAP to prevent evaporation of the agar pad^85^. To follow oocyte mitochondrial abundance during OZT, z-stacks (0.37μm per section) were captured every 5 min for 50 min. Exposure times and laser lower were optimized for each experiment based on fluorescence intensity. To track FUNDC1 puncta on oocyte mitochondria and to track mitochondrial fragmentation events, single slice images were captured every 1.5 s for 8-15 min. The collected images were viewed and analyzed using FIJI software (v1.54f).

Single confocal z-slices of −2 oocytes, −1 oocytes and zygotes from the same focal plane, from the same animal were used as representative images in figures. Fluorescence images were cropped for clarity before displaying in figures. Refer to Supplementary Figures for the uncropped fluorescence images used in main figures (Supplementary Fig. 1) and extended data (Supplementary Fig. 2).

### RNAi treatment

RNAi knockdown was performed by using standard feeding of *Escherichia coli* HT115 expressing the relevant double stranded RNA. RNAi constructs used in this study were obtained from the RNAi libraries constructed by the Ahringer laboratory^86^ and the Vidal laboratory^87^. Bacteria with L4440 or T444T empty vectors were used as relevant controls.

RNAi bacterial strains were cultured in selective media (LB media containing 100 mg/mL ampicillin) for ∼14 h at 37°C. Expression of double stranded RNA was triggered by addition of 1mM IPTG. These bacterial cultures were seeded onto nematode growth medium agar plates treated with 1mM IPTG and 100 mg/mL ampicillin and dried at room temperature overnight before use. All RNAi knockdown experiments were conditional, with target RNAi fed beginning at the L4 stage to avoid developmental defects in the hermaphrodite germline. In every biological replicate, ∼30 synchronized L1 worms were grown on negative control (L4440 or T444T) for 48 h followed by transfer onto relevant RNAi plates for 24 h before examining or imaging mitochondrial degradation at OZT.

### RNAi knockdown efficiency

To measure the knockdown efficiency of *drp-1* and *fzo-1* RNAi, L1 animals expressing endogenous *gfp::drp-1* and *fzo-1:gfp* were grown for 48 h on negative control (L4440) plates before subjecting them to respective RNAi for 24 h at 20°C. Net total fluorescence intensity of GFP::DRP-1 and FZO-1::GFP within the −2 oocyte was measured from background subtracted sum projections (3 z-slices). Knockdown efficiency percentage was calculated using the formula: [Total fluorescence intensity in −2 oocyte of control animal – total fluorescence intensity in −2 oocyte of RNAi treated animal]/ [Total fluorescence intensity in −2 oocyte of control animal] x100. Refer to Supplementary Figure 3 for knockdown efficiency data.

### Quantification of organelle/protein abundances at OZT

Confocal z-stack images (0.37μm per section) of −2 oocytes, −1 oocytes and zygotes expressing TOMM-20::mNG, NDUV-2::mNG, GFP::DRP-1^88^, ATG-9::GFP^89^, mCherry::LGG-2^90,91^, CTNS-1::wrmScarlet^92,93^, FZO-1::GFP or mRb3::FNDC-1 were captured using a x63/1.4NA oil immersion objective. Sum projections were generated from 3-slices after background subtraction in Fiji (15px rolling ball). Regions of interest were manually drawn around the −2 oocytes, −1 oocytes or zygotes and total fluorescence intensity was calculated for the respective cell using the measure tool. For ATG-9 vesicle, autophagosome (mCherry::LGG-2) and lysosome (CTNS-1::wrmScarlet) organelle counts, the “3D object counter” plugin was used over 3 z-slices, with the average fluorescence intensity of the respective marker used to set the threshold for object detection. The same “3D object counter” analysis pipeline was used to calculate the volume of GFP::DRP-1 positive puncta, autophagosome volume or lysosome volume. Between 10-15 animals were quantified per strain in each biological replicate. Three biological replicates were performed per strain.

### Quantification of mitochondrial volume and morphology

Fluorescence images of day 1 adults expressing endogenous NDUV-2::mNG (3 z-series confocal images, 0.37μm per slice) were captured using spinning-disk microscopy. The volume of NDUV-2::mNG-positive mitochondria was quantified using the Fiji macro, “Mitochondria Analyzer”^94^. Images were background subtracted (15px rolling ball), converted to 8-bit and oocytes or zygotes were trimmed after manually tracing outlines around the respective cell before analysis. 3D analysis was performed throughout the acquired z-stack using the settings: Max slope:1.40, gamma 0.90, block size 1.25μm and C-Value 5. ‘Despeckle’, ‘remove outliers’ and ‘fill 3D holes’ were set to be performed post-processing. The same analysis settings on “3D Mitochondria Analyzer” were used for different cells (−2 oocyte vs zygote) and across different strains to accurately measure changes in mitochondrial volume.

Mitochondrial morphology was quantified in the −2 oocyte and zygote using the “3D Mitochondria Analyzer” pipeline and the network parameters “Mean mitochondrial volume (µm³)”, “Mitochondrial count” and “Sphericity” were noted. Between 10-15 animals were quantified per strain in each biological replicate. Three biological replicates were performed per strain.

### Quantification of mitochondrial degradation at OZT

Mitochondrial protein abundance and total mitochondrial volume in the −2 oocyte and zygote were quantified for z-stack images (3 slices, 0.37μm per section) as outlined above. Fold reduction in mitochondria was used to quantify extent of mitophagy during OZT. This was calculated using the formulae: [(Mitochondrial volume in zygote)/ (Mitochondrial volume in −2 oocyte)] and [(Total NDUV-2::mNG fluorescence intensity in zygote)/ (Total NDUV-2::mNG fluorescence intensity in −2 oocyte)]. Fold reduction in mitochondria was quantified for each animal individually and between 10-15 animals were quantified per strain in each biological replicate. An average fold reduction value of ∼0.5 denoted a 2-fold reduction in mitochondrial protein abundance or mitochondrial volume, while a value of ∼1.0 denoted disrupted mitophagy at OZT.

### Staining with mitochondrial dyes

Nonyl Acridine Orange (NAO) was reconstituted in M9 buffer to a concentration of 5μM and added to plates with 8mL NGM to make a final concentration of 125nM NAO. OP50 bacterial food was seeded onto the NAO plates for culturing *C. elegans*. About 30 L4 larval stage animals were picked onto these NAO plates and were allowed to grow in the dark for 24 h at 20°C. Day 1 adults from the NAO plates were then used for imaging mitochondria at OZT. NAO dye photobleached upon acquisition of z-stacks. Thus, single confocal slices were used for quantification of NAO signal in −2 oocytes and zygotes.

### Quantification of mRb3::FNDC-1 localized at OMM and MAMs

Using confocal z-stacks (10-15 z-slices, 0.37μm per section) the total fluorescence intensity of mRb3::FNDC-1 was quantified at the OMM and MAMs after background subtraction (15px rolling ball) in the −2 and −1 oocytes. The total fluorescence intensity within a 1.83 x 1.83 μm (5 x 5 pixels) box at the mitochondrial surface was measured. Ten regions along oocyte mitochondria were sampled and the mean was used for each animal analyzed. This value was used to represent mRb3::FNDC-1 intensity at OMM, denoted by “OMM FUNDC1 fluorescence intensity”. The total fluorescence intensity within each mRb3::FNDC-1 puncta was measured after manually drawing regions of interest around individual mRb3::FNDC-1 puncta. All mRb3::FNDC-1 puncta in the z-stack were included in the sampling and the mean was used.

This value was used to represent mRb3::FNDC-1 intensity at MAM, denoted by “average FUNDC1 puncta fluorescence intensity”. Between 10-15 animals were quantified per treatment in each biological replicate. Three biological replicates were performed per strain.

### Quantification of FUNDC1 puncta co-localization

Fluorescence images of −2 and −1 oocytes in day 1 adults were captured by spinning-disk microscopy. Co-localization analyses of mRb3::FNDC-1 signal and endoplasmic reticulum(GFP::SP12) or mitochondria(NDUV-2::mNG) in and oocytes was performed on single confocal z-sections using the Fiji plugin, “JACoP”. Pearson’s correlation coefficient (r) was calculated from a region of interest drawn around individual mRb3::FNDC-1 puncta. Line scans spanning a 1μm region across individual mRb3::FNDC-1 puncta were performed on fluorescence images of germ cells in strains expressing (mRb3::FNDC-1 with NDUV-2::mNG) and (mRb3::FNDC-1 with GFP::SP12) to further confirm co-localization.

The percentage of mRb3::FNDC-1 that localized on mitochondria in −2 oocytes and −1 oocytes were manually quantified with 10 z-series confocal fluorescence images (0.37μm per section). Pearson’s correlation coefficient (r) was calculated between mitochondria and mRb3::FNDC-1 signal for all FUNDC1 puncta observed in the −2 and −1 oocytes of 10 animals; mRb3::FNDC-1 puncta for which r values were measured to be positive (0.40 – 0.60) were scored as “localized on mitochondria” and the puncta for which r values were below 0.40 were scored as “not localized on mitochondria”. Most non-mitochondrial localized mRb3::FNDC-1 had r values that were negative or close to zero. This validated our manual quantification of percentage of mitochondrial localized mRb3::FNDC-1 puncta in −2 oocytes and −1 oocytes. 30 animals were quantified in total over three biological replicates.

### Quantification of mRb3::FNDC-1 puncta dwell time on mitochondria

Fluorescence images (single z-slice) of −1 oocytes were captured every 1.5 s for 8-15 min in strains expressing NDUV-2::mNG and mRb3::FNDC-1. Dynamic changes in co-localization between mRb3::FNDC-1 and mitochondria in time-lapsing experiments conducted in −1 oocytes were validated by measuring and tracking change in Pearson’s correlation coefficient (r) across time. The timepoint at which Pearson’s correlation coefficient (r) values drop from positive values (indicating co-localization) to zero or negative values was used to mark end of co-localization. The time for which each mRb3::FNDC-1 co-localized on mitochondria was calculated manually, denoted by “dwell time on mitochondria”. Over 30 mRb3::FNDC-1 puncta from 15 animals were tracked and quantified to characterize dynamic behaviors of FUNDC1 puncta in the −1 oocyte.

### Quantification of fragmentation events

Time-lapse images of day 1 adults expressing NDUV-2::mNG and mRb3::FNDC-1 were captured by spinning-disk microscopy. Images were captured every 1.5 s for 8-15 min. Line scans spanning 1μm were performed along thresholded mitochondrial signal before and after fission event to validate break in mitochondrial signal. mRb3::FNDC-1 signal was assessed for at the fission site using line scans. 10 mitochondrial stabilized mRb3::FNDC-1 puncta across four independent experiments were tracked to assess mitochondrial fragmentation.

### Mitochondrial analysis in female animals

Feminization of *C. elegans* was achieved through transgenerational RNAi knockdown of *fem-3*^33^. In each experimental setup, L1 worms were plated onto *fem-3* RNAi plates and 48 h later, L4s were transferred to fresh *fem-3* RNAi plates, and this was repeated for 3 generations at 20°C. Successful feminization was confirmed by the presence or absence of embryos in the animal’s uterus using confocal microscopy. This was performed in the NK2747 (TOMM-20::mNG) and NK2845 (NDUV-2::mNG) strains to assess mitochondrial abundance and morphology. Confocal z-stacks was acquired to examine mitochondria in successfully feminized day 1 adult animals. Care was taken to avoid measurements from older feminized animals displaying oocyte “stacking” phenotype, where numerous oocytes stack against each other over time resulting in oocytes with a columnar shape^95–97^. To avoid confounding factors and assess specifically the effect of absence of sperm on oocyte mitophagy/morphology, we limited sampling to day 1 adults that exhibited the typical 3-5 oocytes along the proximal germline (similar to day 1 hermaphrodites). Hermaphrodites grown on T444T RNAi plates for 3 generations were used as control for mitochondrial analysis.

### UVC exposure

*C. elegans* strains were grown to L4 larval stage. At the time of exposure, 10-15 worms per strain were transferred from plates seeded with OP50, washed once in M9 buffer and plated onto unseeded NGM plates to avoid shielding of UVC treatment by bacteria. The worms were then exposed to 25J/m² UVC radiation of 254nm using a UVC lamp with a built-in UVC sensor (CL-1000 Ultraviolet Crosslinker). UVC radiation is known to induce photodimers^98–101^, which generates cyclobutane pyrimidine dimers in mtDNA that when used as templates for DNA replication, can lead to mutations^98,102^. Following exposure, worms were transferred to NGM plates with OP50 and grown for 24 h at 20°C to allow for DNA mutagenesis.

The criteria to determine the single dosage to use for mtDNA mutagenesis experiment was (i) the dosage caused significant mtDNA lesions at detectable levels, 24 h after exposure (ii) the lesions caused on nuclear DNA (nDNA) from the dosage would be removed within 24 h after exposure through nucleotide excision repair^102^, avoiding nDNA mutations, and (iii) did not have significant effects on organismal fitness in wild-type animals (growth and fecundity). 25J/m² was used as the dosage for mtDNA damage experiments based on previous studies on UVC mediated mtDNA damage in *C. elegans*, which show that this dosage caused substantial mtDNA lesions still detectable 24 h after exposure (∼0.2 mtDNA lesions/10-kb detected immediately post-UVC exposure and ∼0.4 mtDNA lesions/10-kb detected 24hrs post-UVC exposure)^103^. Further, nDNA lesions were significantly reduced 24 h after 25J/m² UVC exposure (more than ∼2-fold decrease)^103^. To ensure 25J/m² did not cause reproductive fitness decline in wild-type animals, 80-90 wild-type L4 animals were treated with 25J/m² UVC radiation and the proportion of gravid adults was counted every 12 h to quantify developmental delays caused by UVC. The number of viable progenies from each animal was counted to quantify reproductive fitness decline. There were no developmental defects (100% of UVC treated wild-type L4 animals reached gravid adulthood at the same rate as untreated L4s) or defects in fecundity (non-significant changes in brood size) upon 25J/m² dosage.

### Mutation accumulation lines

Using the standardized dosage of UVC treatment (25J/m²), mutation accumulation (MA) line experiments were performed in wild-type and *fndc-1 C. elegans*, in control conditions and upon UVC exposure. A single random L4 of each strain was isolated (G0) and allowed to lay eggs.

Once the offspring reached the L4 stage, one worm was randomly picked onto new NGM plate with OP50 to create the G1 subline. 10 different G1 sublines were generated per strain per treatment to start the MA experiment. One individual L4 animal was picked at random and transferred to new plate to be the progenitor for the next generation. In UVC treatment conditions, this L4 progenitor was exposed to 25J/m² UVC once before allowing it to propagate the next generation. This was repeated for an individual L4 in every subsequent generation for the UV MA lines for accumulation of mtDNA mutations. Every subline was subject to such bottlenecking for 10 generations per strain per treatment. All lines were grown and propagated at 20°C. Two previous generations were stored at 16°C, so that if an individual was arrested before adulthood or dead or sterile, a back-up individual could be used to continue MA line. The line was considered extinct if the back-up was also not viable.

### Paraquat, FCCP, and 6PPD-Q treatment

Synchronized L1 animals were grown to L4 larval stage on NGM plates with OP50. At the time of exposure, 10-15 worms were transferred from plates seeded with OP50, washed three times in complete K medium^104^ (2.36 KCl, 3 g NaCl, 5 mg cholesterol, 1 mL 1M CaCl_2_, 1 mL 1M MgSO_4_ in 1 L of water) and grown in wells containing 1 mL complete K medium. Worms were kept from starvation by feeding 50 mg/ml OP50. Oxidative damage was induced through Paraquat treatment^105^, mitochondrial uncoupling through FCCP treatment^106^ and mitochondrial dysfunction through 6PPD-Q treatment^107,108^. Paraquat, FCCP and 6PPD-Q were diluted in DMSO and added to wells with K medium to achieve final dilutions of 75µM^109^, 5µM^110^ and 33µM respectively. The equivalent concentration of DMSO was added to the control wells. Well plates were rocked gently and incubated at 25°C. L4 worms were grown in control and treatment conditions for 24 h and day 1 adults were transferred to slides for confocal imaging.

### *uaDf5* heteroplasmy quantification

mtDNA heteroplasmy via RT-PCR was conducted by adapting previously described protocols^64,111^. For each strain, in each independent experiment, individual germlines were extruded from day 1 adult hermaphrodites and F1 offspring at L1 larvae stage were picked from the same plate and transferred into separate PCR tubes containing lysis buffer. The composition of lysis buffer used was as follows: 60% nuclease-free water, 5% 20mg/µL Proteinase K, and 35% 3.3X lysis buffer. Lysis was set up at ratios of 1 sample/15µL buffer for adult germlines and 1 worm/10µL buffer for L1s. For completion of lysis reaction, samples were stored at −80°C for 10 min, lysed for 1 h at 65°C, and Proteinase K was deactivated at 95°C for 15 min. RT-PCR was then performed on lysates using 25 µl reaction volumes (2µL lysate, 8.5 µL nuclease-free water, 12.5 µL Power SYBR Green Master Mix, and 1 µL of each 100 µM primer). mtDNA heteroplasmy was determined by using two oligo pairs that specifically detected wild-type mtDNA or total mtDNA. Two independent RT-PCR reactions of the same sample were run simultaneously to determine cycle-threshold (Ct) values for wild-type mtDNA and total mtDNA. At least 10 reactions were conducted across three independent experiments, and wild-type and total mtDNA cycle-threshold (Ct) values were used to quantify the relative abundance of *uaDf5* mtDNA in each sample. Reactions where Ct values from the two technical replicates differed by ≥1 cycle were excluded. Mean of Ct values from each technical replicate were reported. *uaDf5* heteroplasmy percentage was calculated as follows:

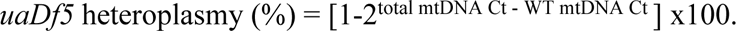

Oligo pairs that specifically detect only wild-type mtDNA and an oligo pair targeting all mtDNA copies were used (Refer Supplementary Table 3):

Wild-type mtDNA Fw: 5’-gtccttgtggaatggttgaatttac-3’ Wild-type mtDNA Rv: 5’-gtacttaatcacgctacagcagc-3’ Total mtDNA Fw (*nd-1*): 5’-agcgtcatttattgggaagaagac-3’ Total mtDNA Rv (*nd-1*): 5’-aagcttgtgctaatcccataaatgt-3’

### Examination of embryonic lethality

To examine embryonic lethality, ten L4 worms were singled onto individual NGM plates seeded with OP50. The worms were then transferred to fresh OP50 plates every 24 h for 6 days. The eggs laid on each plate were counted right after removing the parent worm and added to calculate the total number of eggs laid per animal. 48 h after removal of parent animal, the number of hatched viable progeny was counted along with the number of unhatched dead embryos. The embryonic lethality percentage was calculated using the formula: [Number of dead embryos]/ [Total number of eggs laid] x100. Ten parent animals and all embryos laid by each animal were quantified per strain per treatment.

### Quantification of reproductive fitness

Ten L4 animals per strain were singled onto individual NGM plates seeded with OP50. When worms began laying eggs (day 1 adulthood), they were transferred to new plates every 24 h for 6 days. 48 h after removal of parent animal, the number of live progeny was counted. The number of viable progeny counted per plate were added to calculate total brood size per animal. A minimum of 10 worms were used for quantification per strain per treatment.

### Mating experiments

Ten males expressing *Itls44* [*pie-1p::mCherry::PH(PLC1delta1)*], marking germline and embryonic plasma membrane, were placed on an NGM agar plate with a small OP50 bacterial lawn. Two *fndc-1*(*rny15*) L4 stage hermaphrodites from *fndc-1* (UV) MA lines (G4) were transferred individually to the center of the lawn. Efficient mating was confirmed by assessment of red fluorescence (mCherry) in progeny through confocal imaging. 10 separate crosses were set up with 2 hermaphrodites from each of the 10 *fndc-1* (UV) MA lines.

### Mitochondrial respiration

Day 1 adult *C. elegans* from wild-type MA lines, wild-type (UV) MA lines, *fndc-1* MA lines, and *fndc-1* (UV) MA lines were picked off NGM plates with OP50 and the Seahorse Extracellular Flux Bioanalyzer was used to quantify respiration parameters as described in previous studies^112^. Confocal images of the plate were acquired to count the number of individual animals per well, such that Oxygen Consumption Rate (OCR) measurements were normalized per individual worm. 5-10 adult animals from each of the 10 MA lines were used per strain per condition across four independent experiments. Means of technical replicates per plate were reported. Effect of UV treatment across the 2 different strains were compared using two-way ANOVAs.

### Statistical analysis

All experiments were performed in replicates of three. Figure legends include details about sample size and statistical analysis used. Statistical analyses were performed in GraphPad Prism 10 and GraphPad 2×2 contingency table. Normality of data distribution was assessed using the Kolmogorov-Smirnov test. Data derived from two different conditions/cells were compared using two-tailed Student’s *t*-test (Gaussian distribution), Mann-Whitney *U* test or Wilcoxon test (non-Gaussian distribution). For comparisons of three or more datasets, one-way ANOVA followed by Tukey’s multiple comparison test (Gaussian distribution) or Kruskal-Wallis test followed by Dunn’s multiple comparisons test (non-Gaussian distribution) were performed.

Categorical data was compared using two-tailed Fisher’s exact test (RNAi screen in Supplementary Table 1). Data were considered statistically different at P<0.05. All data are generally expressed as mean ± s.d. except when mentioned to be mean ± s.e.m. in figure legend.

## Data availability

All data are available in the main text and supplementary materials. Illustrations were prepared using the programs Procreate (https://procreate.com/procreate) and Adobe Illustrator v.29.2.1 (https://www.adobe.com/products/illustrator.html). Supplementary Videos were edited using Adobe Premiere Pro (https://www.adobe.com/products/premiere.html). Confocal images were processed using ImageJ (v.2.1.4.7). Statistical analyses were performed using GraphPad Prism (v.10). Uncropped versions of confocal images used in main and extended data figures are provided in Supplementary Fig. 1 and 2.

## Code availability

No custom code was developed or used in this study.

---

**Extended Data Fig. 1:**
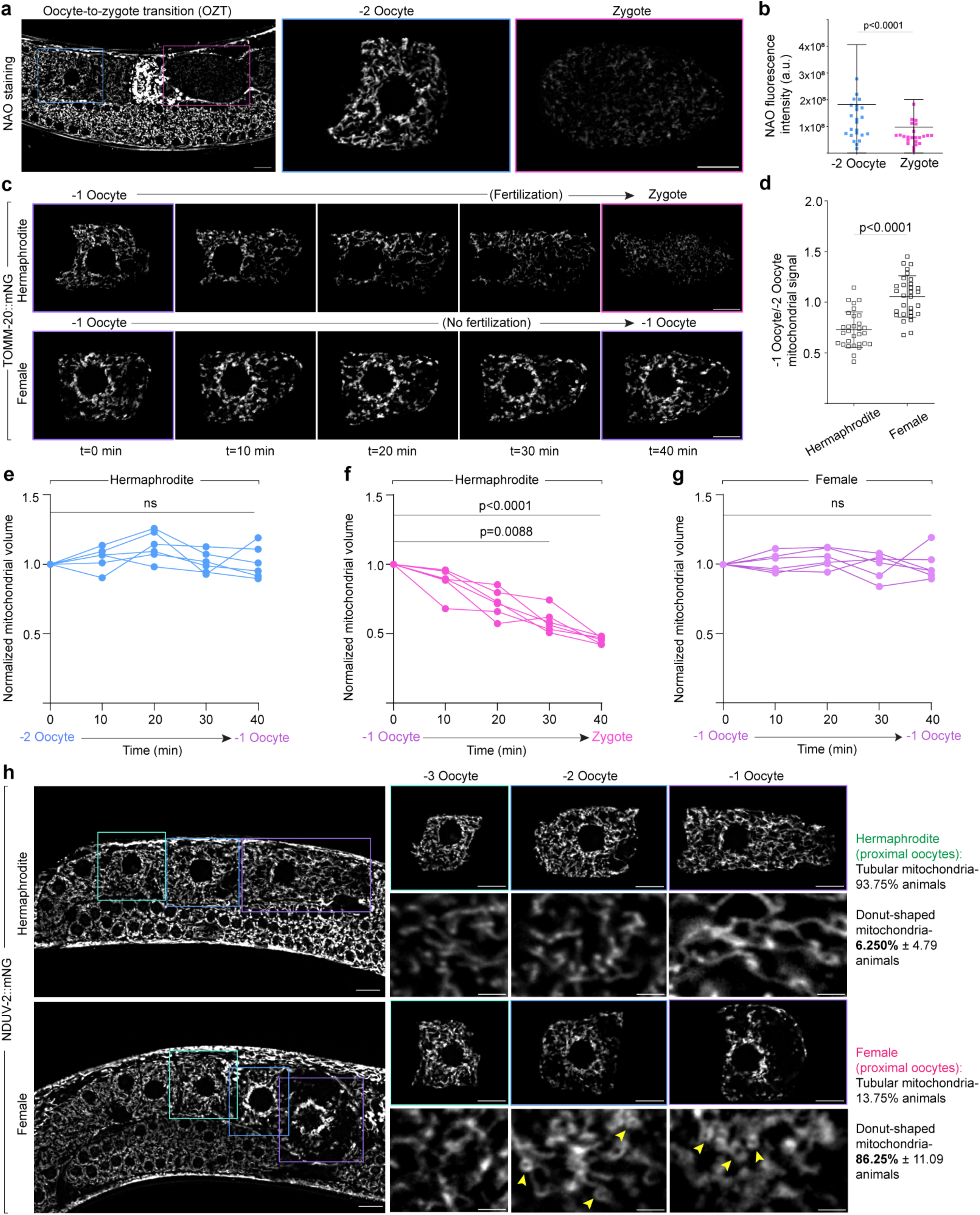
Mitochondrial reduction during OZT is developmentally programmed. **a**, Confocal fluorescence images of mitochondria during *C. elegans* OZT. Mitochondria are labelled using Nonyl acridine orange (NAO), a green fluorescent mitochondrial dye. Blue boxes indicate −2 oocyte and pink boxes zygote. −2 oocytes and zygotes enclosed in boxed regions are magnified. Scale bars, 10µm. **b**, Total fluorescence intensity of NAO in −2 oocytes and zygotes. The data represent the mean ± s.d. (n = 30 animals) from three biological replicates. P value using a Mann-Whitney test. **c**, Time-lapse images showing changes in mitochondrial abundance over 40 min in −1 oocytes of a hermaphrodite vs feminized animal. Fluorescence images show mitochondrial reduction in −1 oocyte during transition to zygote in hermaphrodite (top), compared to unchanged mitochondrial abundance in arrested −1 oocytes in feminized animals (bottom). **d**, Quantification of mitochondrial reduction from −2 to −1 oocyte in hermaphrodite vs feminized animals. Data represent ratio of total NDUV-2::mNG fluorescence intensity in −1 oocyte to that in −2 oocyte compared between hermaphrodites and feminized animals. The data represent the mean ± s.d. (n = 30 animals each condition) from three biological replicates. P value was using a two-tailed Student’s *t*-test. **e**,**f**,**g**, Changes to mitochondrial volume during the hermaphrodite −2 oocyte transitioning to −1 oocyte (**e**), during hermaphrodite OZT (**f**) and in the −1 oocyte of feminized animals (**g**). Mitochondrial volume at each time point is normalized to mitochondrial volume at t = 0. The data represent six independent experiments for each condition. P values using a Kruskal-Wallis test followed by a Dunn’s multiple comparisons test. **h**, Fluorescence images of mitochondria (NDUV-2::mNG) in oocytes of hermaphrodite and feminized *C. elegans*. Green boxes indicate −3 oocytes, blue boxes −2 oocytes and purple boxes - 1 oocytes. Oocytes enclosed in boxed regions are magnified. Yellow arrows indicate donut-shaped mitochondria. Scale bars, 10µm. Right: Data represent mean ± s.d. (n = 30 animals per condition) from three biological replicates.

**Extended Data Fig. 2:**
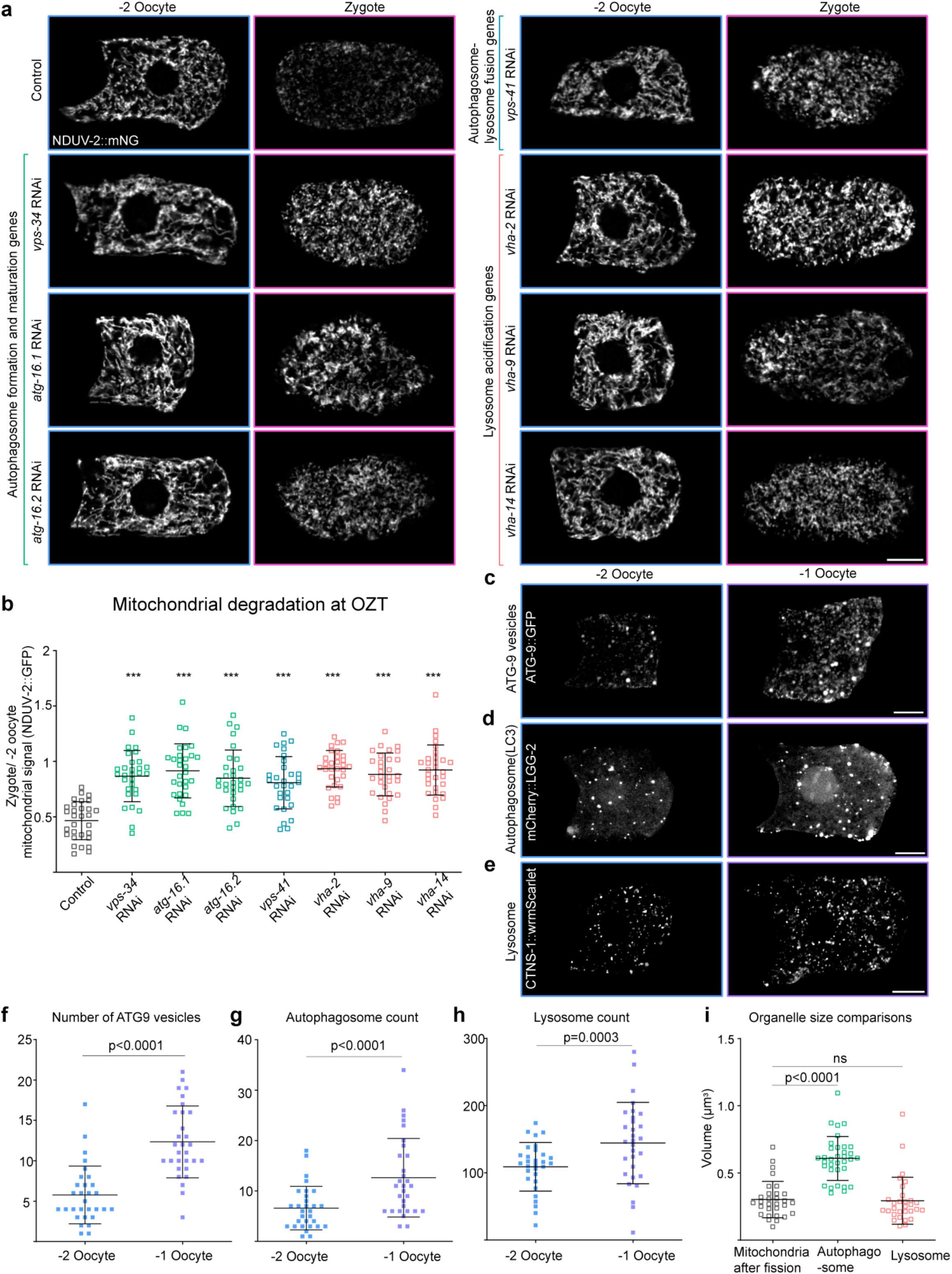
Macroautophagy activation is necessary for mitochondrial degradation during OZT. **a**, Confocal fluorescence images of mitochondria (NDUV-2::mNG) in the −2 oocyte and zygote of empty vector control (L4440) compared to animals treated with RNAi targeting genes involved in macroautophagy. Scale bar, 10µm. **b**, Quantification of mitochondrial reduction during OZT in controls vs RNAi-treated animals. Data represent ratio of total NDUV-2::mNG fluorescence intensity in zygote to that in the −2 oocyte for each animal, compared between control and RNAi knockdown conditions represented in (**a**). The data represent the mean ± s.d. (n = 30 animals each condition) from three biological replicates. ***P<0.0001 from one-way ANOVA followed by Tukey’s multiple comparison test. **c**, Endogenous ATG-9::GFP distribution in −2 oocyte and −1 oocyte. Scale bar, 10µm. **d,** Germline-specific LC3 (mCherry::LGG-2) distribution in the −2 oocyte and −1 oocyte. Scale bar, 10µm. **e,** Confocal fluorescence images of lysosomes (visualized using CTNS-1::wrmScarlet) in - 2 oocyte and −1 oocyte. Scale bars, 10µm. **f**,**g**,**h**, Number of ATG-9 vesicles (**f**), autophagosomes (mCherry::LGG-2) (**g**), and lysosomes (CTNS-1::wrmScarlet) (**h**) in −2 oocytes and −1 oocytes. Data represent mean ± s.d. (n = 30 animals per strain) from three biological replicates. P values using a Wilcoxon test (**f**,**g**) and a two-tailed paired Student’s *t*-test (**h**). **i**, Quantification of mean volume (µm³) of individual fragmented mitochondrion, autophagosome and lysosome in the −1 oocyte. Data represent mean ± s.d. (n = 30 animals per strain) from three biological replicates. P values using a Kruskal-Wallis test followed by a Dunn’s multiple comparisons test. Supplementary Fig.2 shows uncropped fluorescence images.

**Extended Data Fig. 3:**
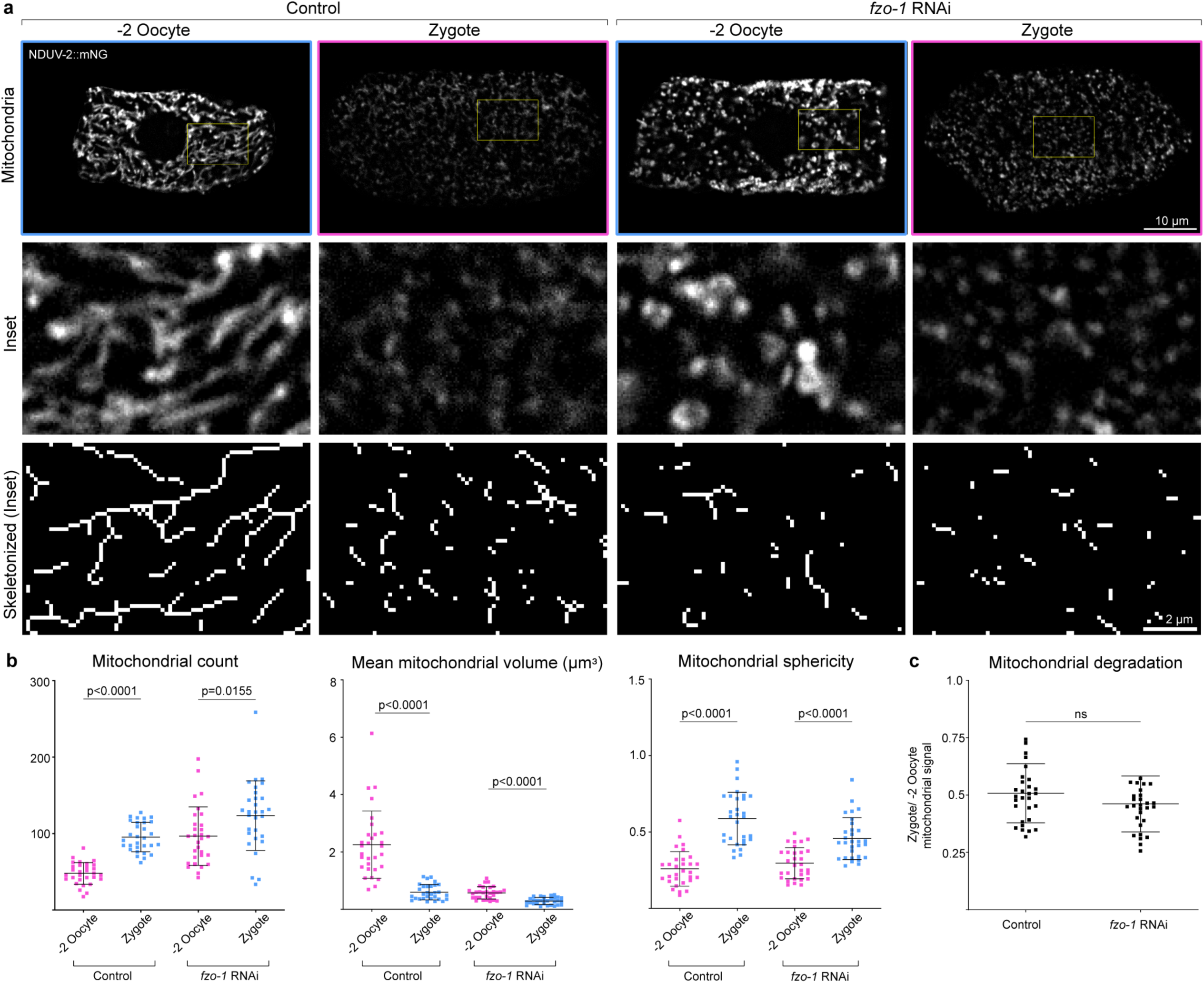
Mitochondrial fragmentation is not sufficient to regulate timing and extent of MOZT. **a**, Top: Confocal fluorescence images showing the change in mitochondrial morphology (visualized with NDUV-2::mNG) between −2 oocyte and zygote in an empty vector control compared to a *fzo-1* RNAi-treated animal. Scale bar, 10µm. Boxed regions are magnified under original images. Bottom: Skeletonized images of the mitochondria shown in middle panels. Scale bar, 2µm. **b**, Quantification of mitochondrial network parameters (NDUV-2::mNG) compared between −2 oocytes and zygotes in control and *fzo-1* RNAi-treated animals. Data represent mean ± s.d. (n = 30 animals per condition) from three biological replicates. P values using two-tailed Student’s *t*-tests and Mann-Whitney tests, depending on normality of distribution. **c**, Quantification of MOZT in control vs *fzo-1* RNAi treated animals. Data represent ratio of total NDUV-2::mNG fluorescence intensity in zygote to that in the −2 oocyte for each animal. The data represent the mean ± s.d. (n = 30 animals each condition) from three biological replicates. P value using a two-tailed Student’s *t*-test.

**Extended Data Fig. 4:**
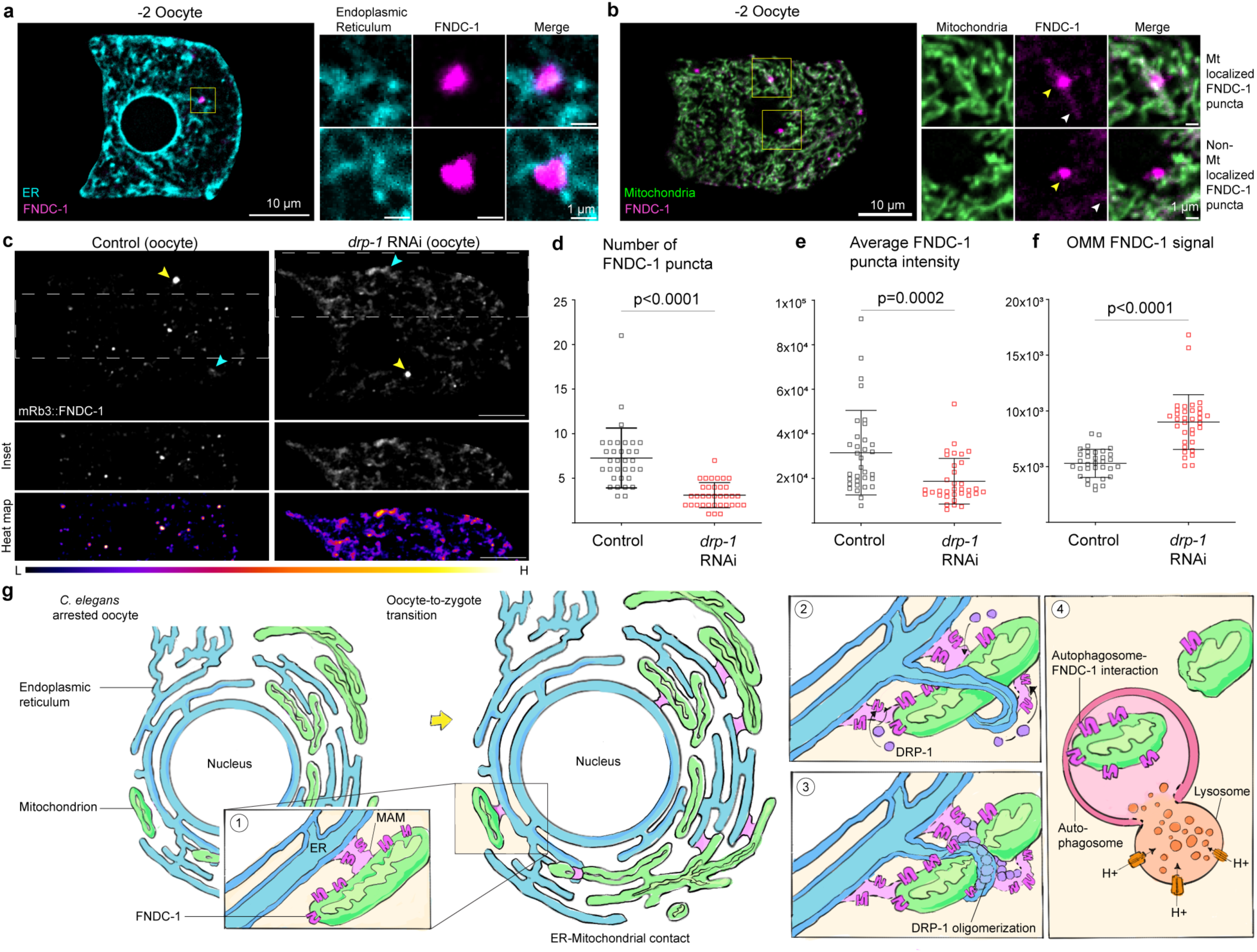
FUNDC1 puncta are associated with ER and mitochondria, marking mitochondria associated membranes. **a,** Left: Merged confocal fluorescence images of the endoplasmic reticulum (visualized with SP12::GFP) and mRb3::FNDC-1 in the −2 oocyte. Scale bar, 10µm. Right: Boxed region is magnified to the right along with another example of FUNDC1 puncta associated with the ER. Pearson’s correlation coefficient (r) was calculated to be 0.4115 ± 0.1762 (mean ± s.d. from n = 15 animals) between FUNDC1 puncta and the ER, showing positive overlap. Scale bar, 1µm. **b,** Left: Merged confocal fluorescence images of the mitochondria (NDUV-2::mNG) and mRb3::FNDC-1 in the −2 oocyte. Scale bar, 10µm. Right: Boxed regions are magnified to the right. Yellow arrows indicate FUNDC1 puncta. White arrows indicate OMM localized FUNDC1 signal. Top right panel shows an example of FUNDC1 puncta associated with the mitochondrial network. Pearson’s correlation coefficient (r) was calculated to be 0.5754 ± 0.1398 (mean ± s.d. calculated from n = 20 animals) for FUNDC1 puncta localized on mitochondria, showing positive overlap. Bottom right panel shows example of FUNDC1 puncta adjacent but not overlapping the mitochondrial network. r = −0.1325 ± 0.1613 (mean ± s.d. calculated from n = 20 animals) for non-mitochondria localized FUNDC1 puncta. Scale bar, 1µm. **c**, Top: Endogenous mRb3::FNDC-1 distribution in the −2 oocyte of control animal compared to a *drp-1* RNAi-treated animal. Yellow and cyan arrows indicate MAM and OMM localized mRb3::FNDC-1 signal, respectively. Region within dashed lines is segmented below. Bottom: Spectral fluorescence-intensity map of segmented region, which displays the minimum (L, low) and maximum (H, high) pixel value of original image. Scale bars, 10µm. **d**,**e**,**f**, Number (**d**), mean fluorescence intensity (**e**) of mRb3::FNDC-1 puncta, and mean fluorescence intensity of OMM localized mRb3::FNDC-1 signal (**f**), in −2 oocytes of controls compared to *drp-1* RNAi-treated animals. Data represent mean ± s.d. (n = 30 animals per condition) from three biological replicates. P values using a Mann-Whitney test. **g**, Proposed model of FUNDC1 localizations and function during *C. elegans* OZT.

**Extended Data Fig. 5:**
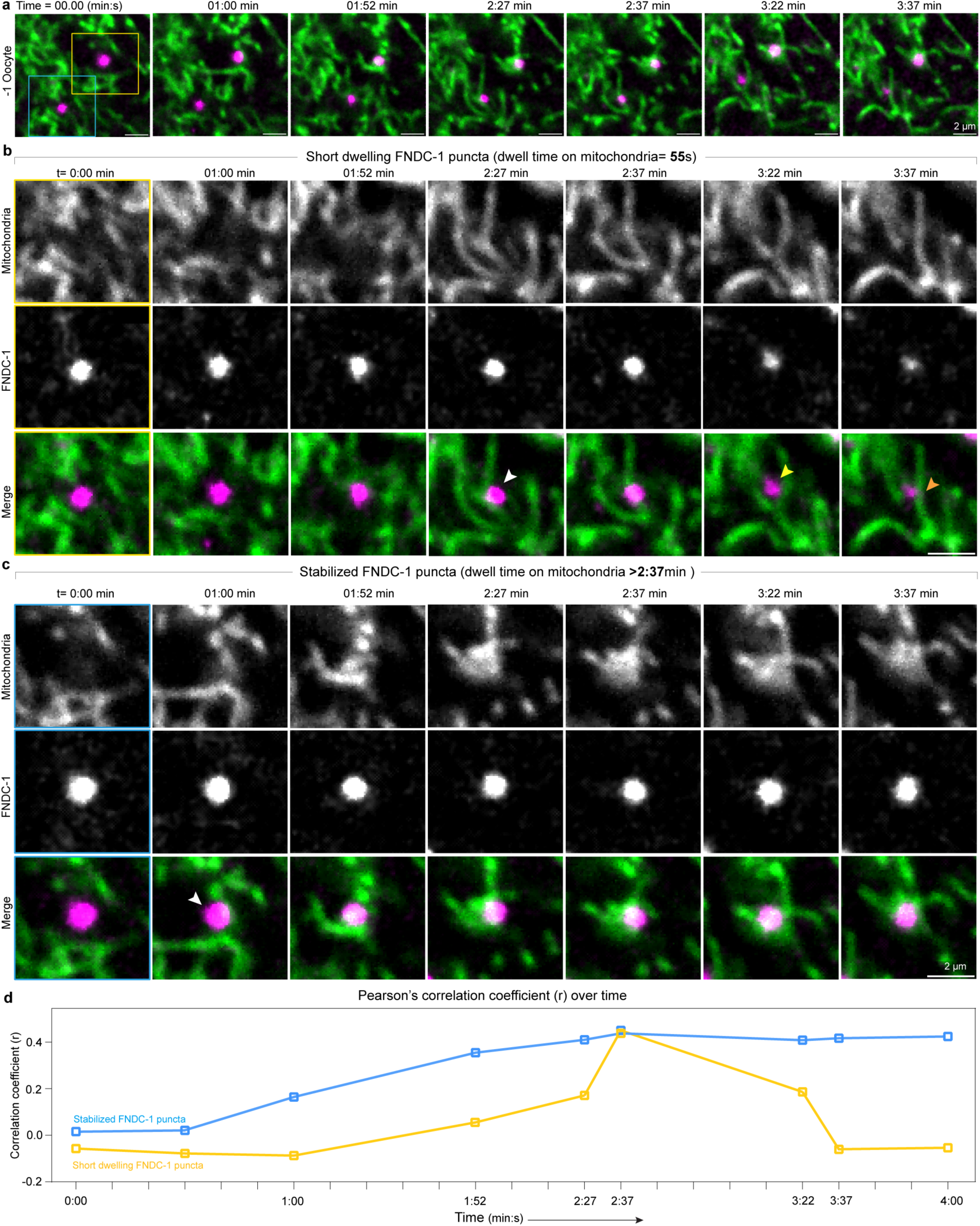
FUNDC1 punctae exhibit two distinct behaviors on mitochondria in the −1 oocyte. **a**, Time-lapse images showing dynamic associations of FUNDC1 punctae with mitochondria in the −1 oocyte. FUNDC1 punctae observed are not co-localized on mitochondria at t = 0 min, but both gradually localizes onto mitochondria. Scale bars, 2µm. FUNDC1 puncta enclosed in yellow and blue boxes are magnified in **b** and **c**. **b**, Time-lapse images show FUNDC1 puncta start localizing on mitochondrial surface (NDUV-2::mNG) around t = 2:27 min, fully co-localizes on mitochondria 10 s later, stays associated with mitochondria for 45 s, after which it moves away from mitochondria. White arrow indicates the start of FUNDC1 co-localization with mitochondria. Yellow arrow indicates the end of FUNDC1 co-localization with mitochondria. Orange arrow indicates intact mitochondria after FUNDC1 puncta is no longer mitochondrial localized. Scale bar, 2µm. **c**, Time-lapse images show FUNDC1 puncta start localizing on mitochondrial surface around t = 1:00 min and stays fully associated with mitochondria for the rest of the time-lapsing experiment (2:37 min). This data represents behavior of a subset of FUNDC1 puncta that stabilize on mitochondria for an extended timespan (4 - 8.6 min). White arrow indicates the start of FUNDC1 co-localization with mitochondria. Scale bar, 2µm. Supplementary video 2 shows the time-lapse data from which **a**, **b,** and **c** are derived. Data represent FUNDC1 puncta dynamics observed in −1 oocytes of n = 35 animals from 8 independent experiments. **d**, Pearson’s correlation coefficient (r) plotted over time, measuring colocalization of FUNDC1 puncta and mitochondria for each frame shown in time-lapses **b** and **c**.

**Extended Data Fig. 6:**
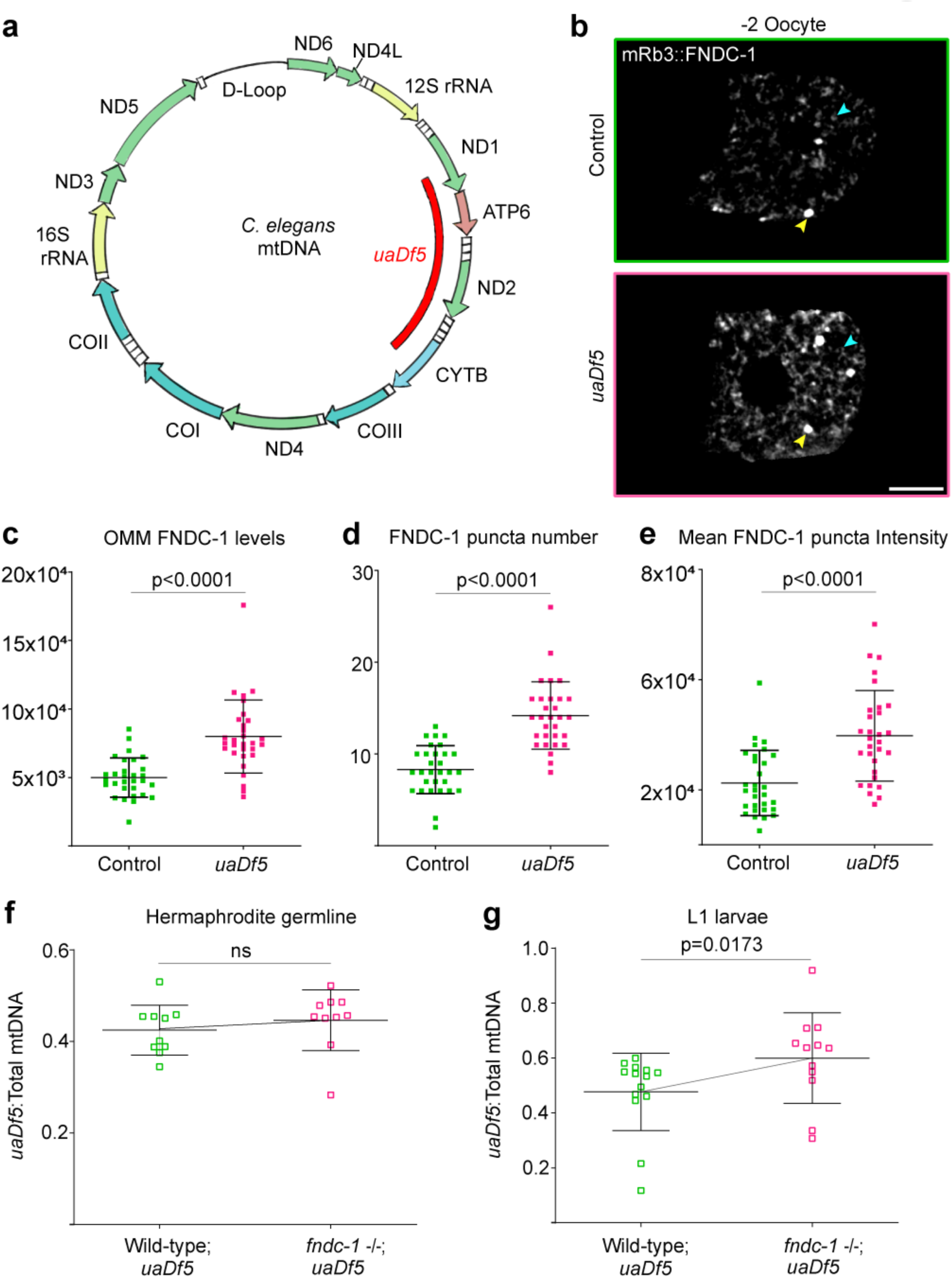
FUNDC1 mitophagy reduces the load of deleterious mtDNA inherited by *C. elegans* progeny. **a**, Schematic of *C. elegans* mtDNA specifying *uaDf5* deletion region (red bar). Arrows show mitochondrial genes encoding proteins and rRNA and their orientation. White boxes show tRNA-encoding genes. Arrows denote genes encoding components of complex I (green), complex III (blue), complex IV (cyan), ATP synthase (brown) and rRNAs (yellow) respectively. Blank line denotes putative D-loop region. **b**, Endogenous mRb3::FNDC-1 distribution in the −2 oocytes of a wild-type control animal compared to an animal harboring the *uaDf5* mtDNA deletion. Blue and yellow arrows indicate OMM and MAM localized mRb3::FNDC-1 signal, respectively. Scale bar, 10µm. **c**,**d**,**e**, Mean fluorescence intensity of OMM localized mRb3::FNDC-1 (**c**), number (**d**), and mean fluorescence intensity (**e**) of mRb3::FNDC-1 puncta in −2 oocytes of controls compared to UVC treated animals. Data represent mean ± s.d. (n = 30 animals per condition) from three biological replicates. P values using Mann-Whitney tests (**c**,**e**) and a two-tailed Student’s *t*-test (**d**). **f**, Comparison of *uaDf5* heteroplasmy between adult germlines of wild-type and *fndc-1* mutant backgrounds containing *uaDf5* deletion. Data represents the ratio of mtDNA copies harboring *uaDf5* to total mtDNA copies (*uaDf5*: total mtDNA) in germlines of day one adult wild-type and *fndc-1* mutants. Data represent mean ± s.d. (n = 10 germlines per strain) from three biological replicates. P value using a Mann-Whitney test. **g**, Comparison of *uaDf5* heteroplasmy between L1 progeny of wild-type and *fndc-1* mutant parents containing *uaDf5* deletion. Data represents the ratio of mtDNA copies harboring *uaDf5* to total mtDNA copies (*uaDf5*: total mtDNA) in L1 progeny of wild-type and *fndc-1* mutants. Data represent mean ± s.d. (n = 14, 12 L1 larvae for wild-type and *fndc-1* mutant backgrounds, respectively) from three biological replicates. P value using a Mann-Whitney test.

**Extended Data Fig. 7:**
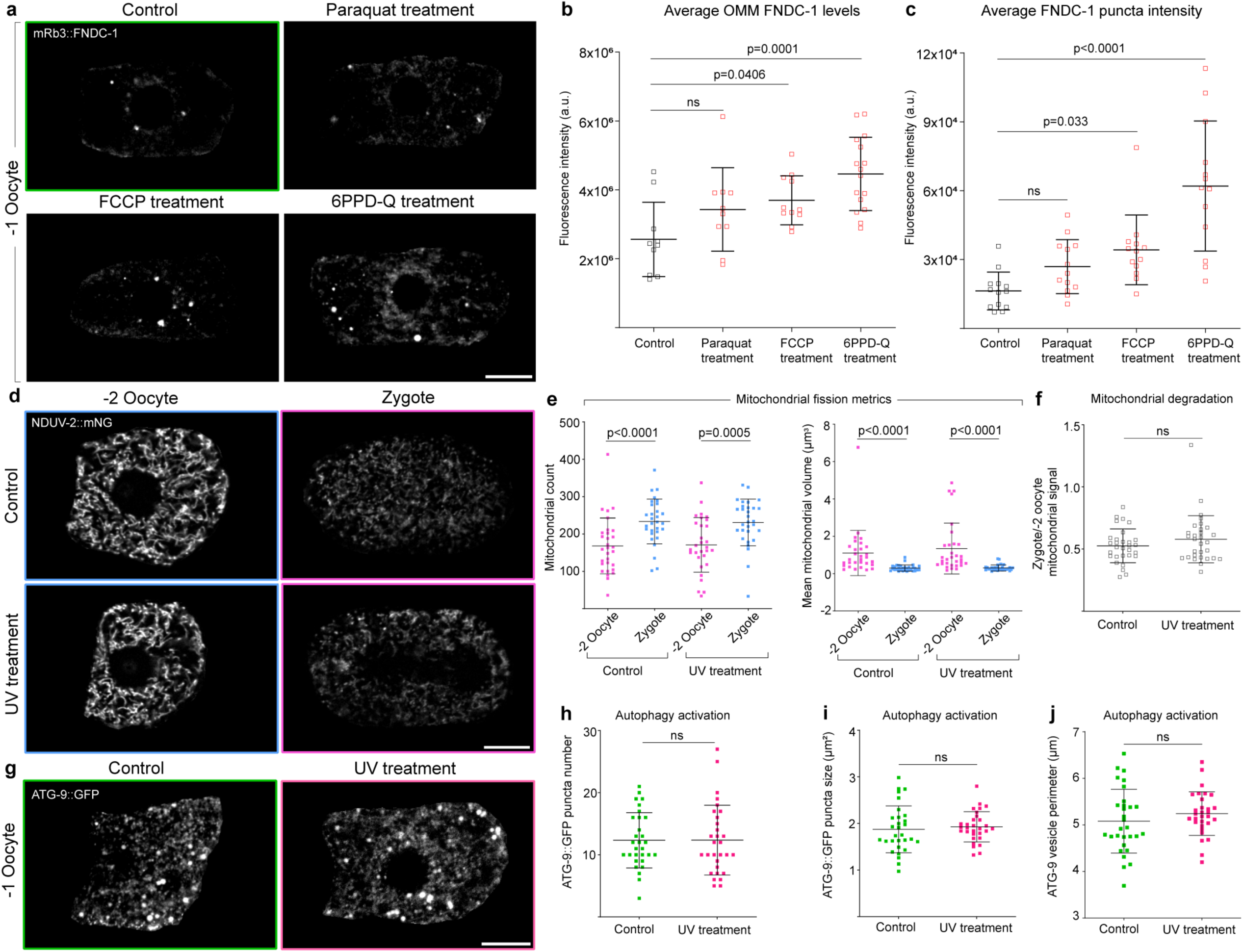
Upon mitochondrial stress, oocytes upregulate FUNDC1 levels, but not mitophagy or mitochondrial fragmentation. **a**, Endogenous mRb3::FNDC-1 distribution in oocytes of a control animal compared to Paraquat, FCCP, and 6PPD-Q treated animals. Scale bar, 10µm. **b**,**c**, Mean fluorescence intensity of OMM localized (**b**) and MAM localized (**c**) mRb3::FNDC-1 in oocytes of control animals compared to animals treated with Paraquat, FCCP, and 6PPD-Q. Data represent mean ± s.d. (n = 10 control, 10 Paraquat treated animals, 11 FCCP treated animals and 16 6PPDQ treated animals) from two independent experiments. P values using a one-way ANOVA followed by a Tukey’s multiple comparison test. **d**, Confocal fluorescence images showing the change in mitochondrial abundance and morphology (visualized with NDUV-2::mNG) between the −2 oocyte and zygote in a control animal compared to a UVC treated animal. Scale bar, 2µm. **e**, Quantification of mitochondrial network parameters compared between −2 oocytes and zygotes in control and UVC treated animals. Data represent mean ± s.d. (n = 30 animals per condition) from three biological replicates. P values using two-tailed Student’s *t*-tests (mitochondrial count) and Mann-Whitney test (mitochondrial volume). **f**, Quantification of MOZT in control vs UVC treated animals. Data represent ratio of total NDUV-2::mNG fluorescence intensity in zygote to that in the −2 oocyte for each animal. The data represent the mean ± s.d. (n = 30 animals each condition) from three biological replicates. P value using a two-tailed Student’s *t*-test. **g**, Distribution of endogenous ATG-9::GFP in −1 oocytes of a control and UVC treated animal. Scale bar, 10µm. **h**,**i**,**j**, Quantification of number (**h**), size (surface area, µm²), (**i**) and perimeter (**j**) of ATG-9 vesicles in −1 oocytes of control vs UVC treated animals. Data represent mean ± s.d. (n = 30 animals per strain) from three biological replicates. P values were calculated from a Mann-Whitney test (**h**) and a two-tailed Student’s *t*-tests (**i,j**).

**Extended Data Fig. 8:**
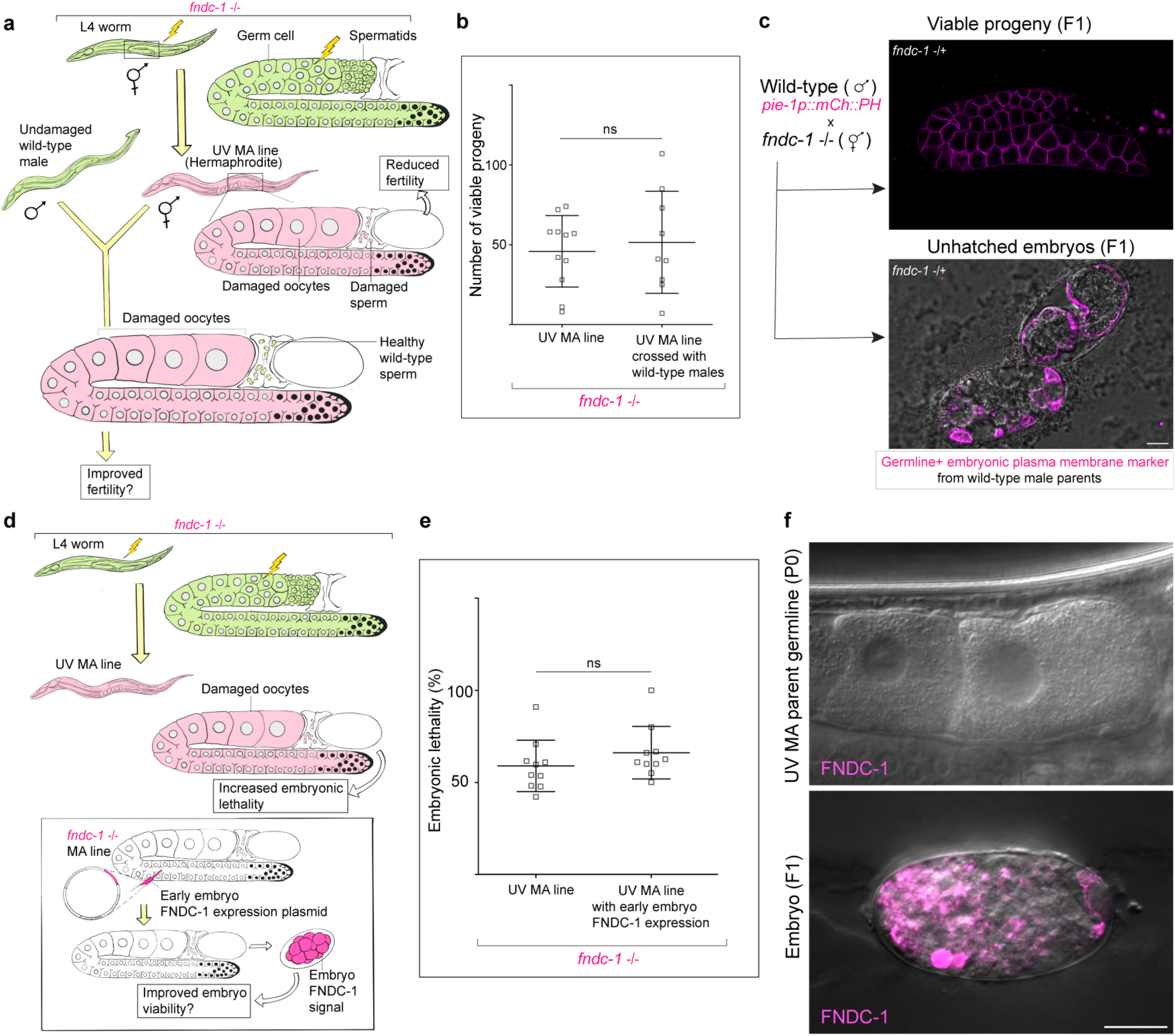
Reproductive decline in *fndc-1* mutant lines upon mtDNA damage is strictly maternal effect. **a**, Schematic of experimental design to rescue sperm health in *fndc-1* UV MA lines. Green animals denote untreated worms, and magenta animals denote worms with mtDNA damage. **b**, Quantification of reproductive fitness (brood size) of self-fertilizing hermaphrodites from *fndc-1* UV MA lines (G4) compared to that of *fndc-1* UV MA line (G4) hermaphrodites crossed with untreated wild-type males. Data represent mean ± s.d. from (n = 10 animals per condition) from three biological replicates. P values using a two-tailed Student’s *t*-test. **c**, Verification of successful crossing between wild-type males and hermaphrodites from *fndc-1* UV MA lines (G4). Merged confocal images showing germline and embryos with plasma membrane fluorescence signal (mCherry::PH, magenta; DIC, grey) among F1 progeny resulting from the cross. Scale bar, 10µm. **d**, Schematic of experimental design to rescue embryonic expression of FUNDC1 in *fndc-1* UV MA lines (G4). Green animals denote untreated conditions, and magenta animals denote mtDNA mutagenized worms. **e**, Embryonic lethality among F1 progeny of parents from the *fndc-1* UV MA lines (G4) compared to F1 progeny of parents (from the same lines) injected with FNDC-1 rescuing embryo expression plasmid [*pie-1p::mRb3::FNDC-1::tbb-2* 3’UTR]. Data represent mean ± s.d. (n = 10 animals per condition) from three biological replicates. P values using a Mann-Whitney test. **f**, Merged confocal fluorescence images of mRb3::FNDC-1 (FNDC-1, magenta; DIC, grey) in oocytes of a *fndc-1* UV MA (G4) parent injected with embryo rescue plasmid and the resulting F1 embryo. Magenta signal in the F1 embryo, but not the parent oocytes represent embryo specific rescue of mRb3::FNDC-1 expression. Scale bar, 10µm.

**Extended Data Fig. 9:**
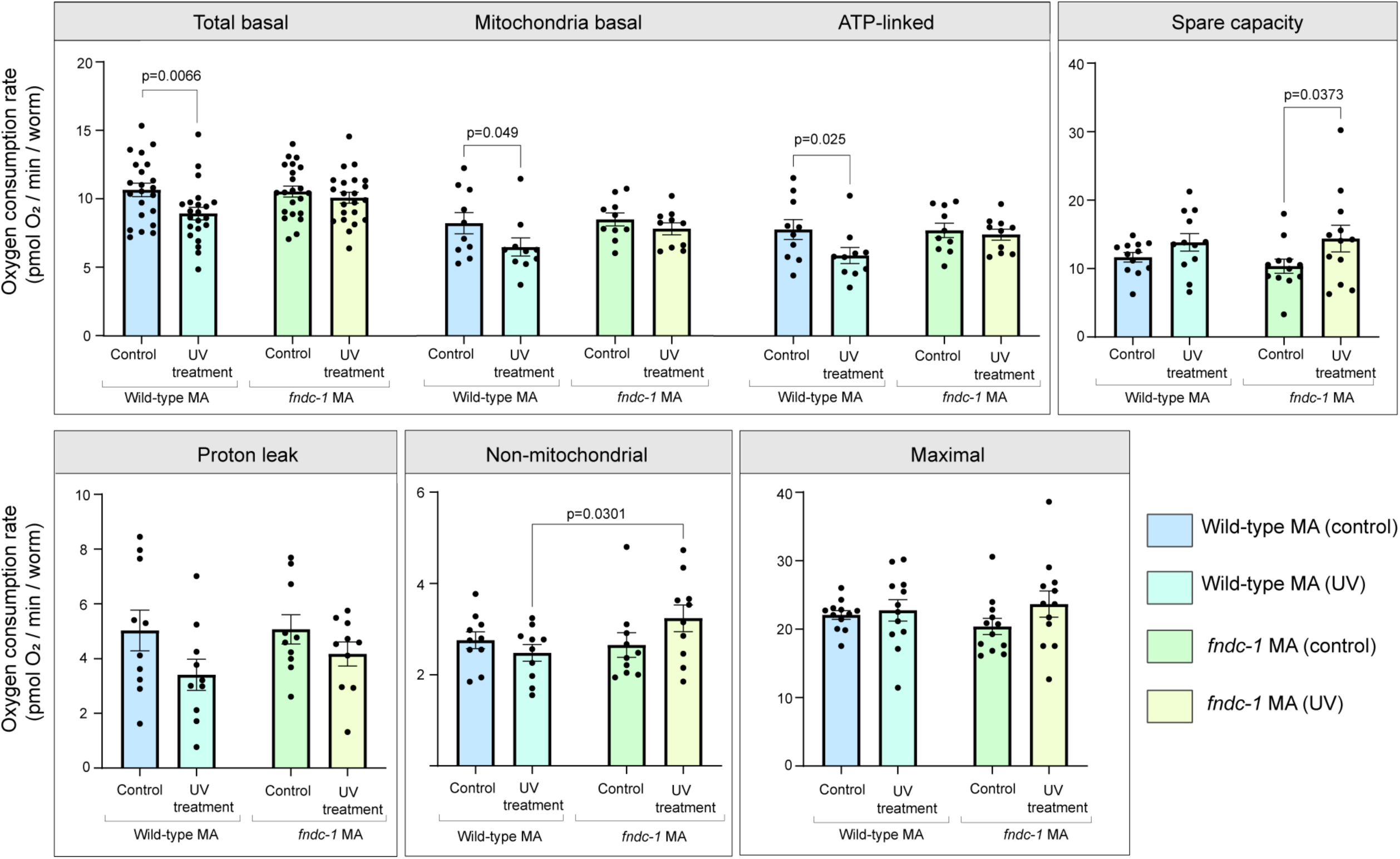
Effect of UV treatment on mitochondrial function in wild-type and *fndc-1* MA lines. Respiration parameters of day 1 adults from the fourth generation (G4) of wild-type control MA lines, wild-type UV MA lines, *fndc-1* control MA lines, and *fndc-1* UV MA lines. Oxygen consumption rate (OCR) measurements were normalized per worm. Data represented are mean OCR/worm value ± s.e.m (n = 10 MA lines) from four independent experiments. Among the UV MA lines, the only respiration parameter that was significantly affected depending on genotype was non-mitochondrial OCR. *fndc-1* UV MA lines exhibited ∼13.07% increased non-mitochondrial OCR compared to wild-type (UV) MA lines (P=0.030). Total basal, mitochondrial respiration, and ATP-linked OCR were significantly diminished (P= 0.0066, P= 0.049 and P=0.025 respectively) in wild-type UV MA lines compared to wild-type control MA lines, showing that mtDNA damage accumulation over multiple generations affected mitochondrial function. UV treatment did not significantly affect respiratory parameters in the surviving animals of *fndc-1* MA lines. Notably, these metrics are reflective of only the ∼45% of animals that successfully hatched and the fraction of the hatched animals that develop to adulthood in each *fndc-1* UV MA line (survivorship bias). In the wild-type UV MA lines there was no embryonic lethality or developmental defects, allowing even sampling. P values using two-way ANOVAs.

**Extended Data Fig. 10:**
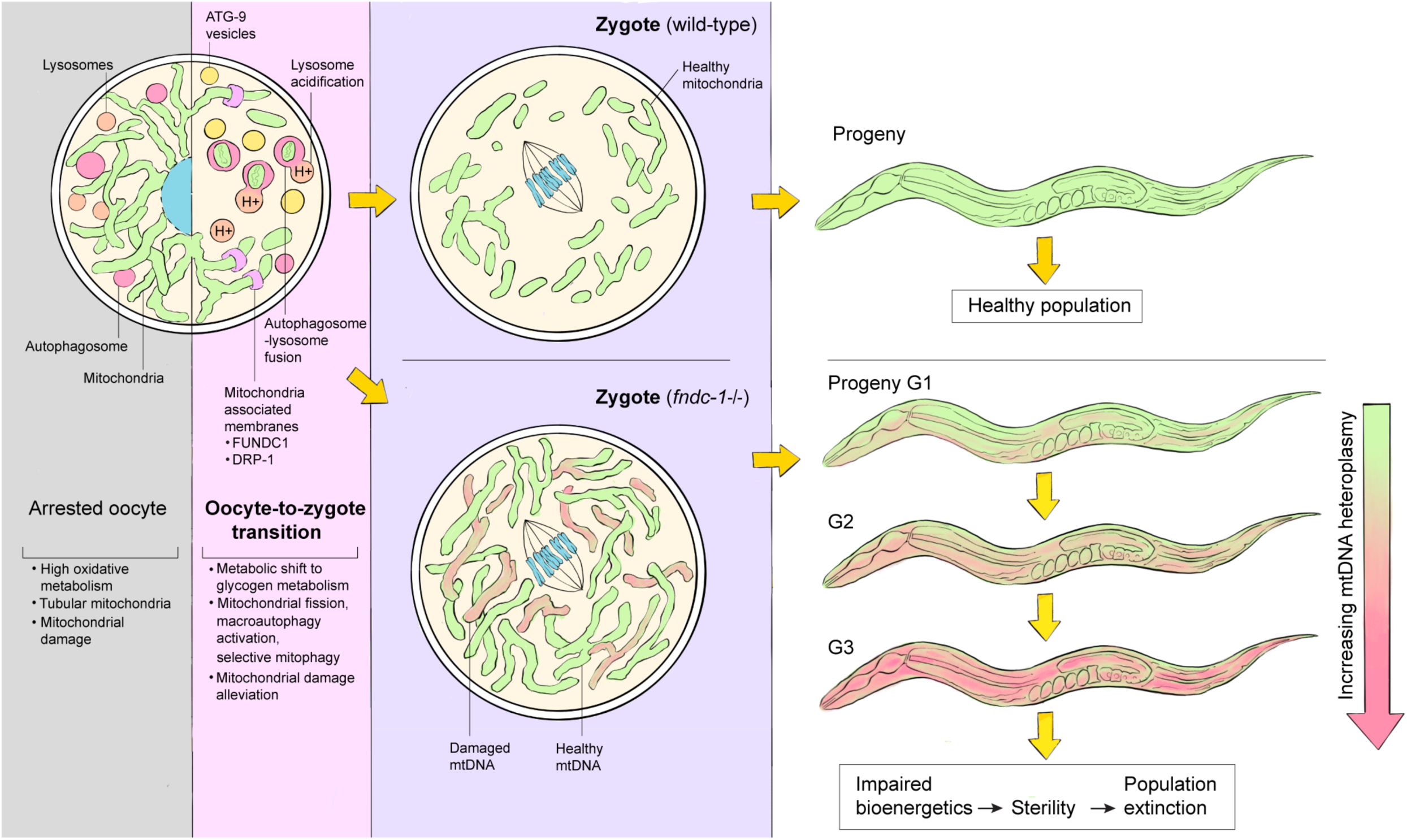
FUNDC-1 mediated programmed mitophagy at oocyte-to-zygote transition (MOZT) promotes mtDNA health, bioenergetics and fecundity in progeny and thereby preserves species continuity.

**Supplementary Video 1**

Mitochondrial fragmentation events during *C. elegans* OZT; FUNDC1 puncta mark sites of mitochondrial fragmentation.

**Supplementary Video 2**

Dynamic interactions of FUNDC1 puncta with −1 oocyte mitochondria.

**Supplementary Fig. 1:**
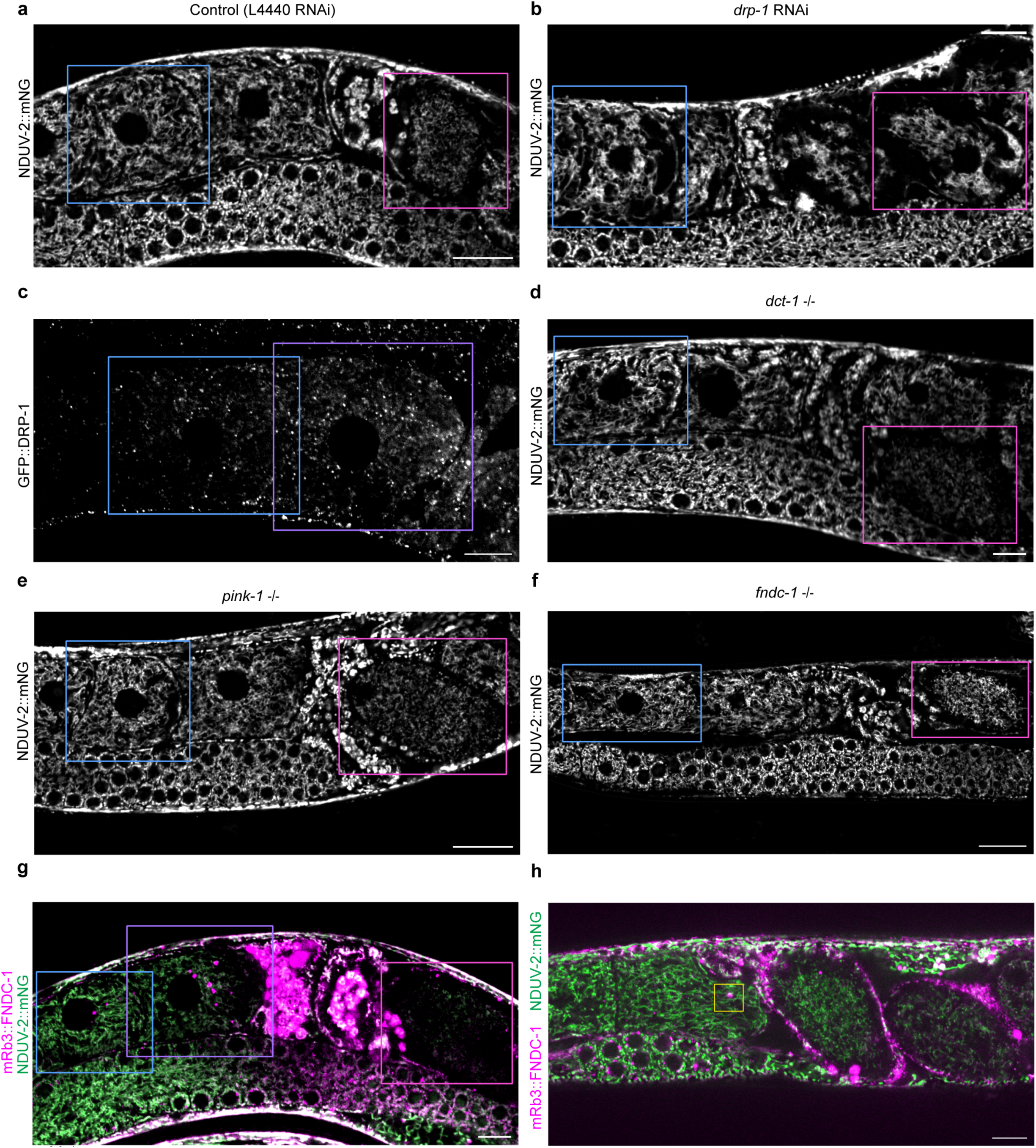
Uncropped confocal fluorescence images from main figures 2, 3 and 4. **a**,**b**, Shown in Fig. 2a. Uncropped fluorescence images showing mitochondria (visualized with NDUV-2::mNG) in the −2 oocyte, −1 oocyte and zygote in an empty vector control (L4440) (**a**) compared to a *drp-1* RNAi-treated animal (**b**). **c**, Shown in Fig. 2e. Uncropped fluorescence images showing endogenous GFP::DRP-1 distribution in the −2 oocyte and −1 oocyte. **d**,**e**,**f**, Shown in Fig. 3a. Uncropped fluorescence images showing mitochondria (NDUV-2::mNG) in the −2 oocyte, −1 oocyte and zygote in null mutants of *dct-1*(*luc194*) (**d**), *pink-1*(*tm1779*) (**e**), and *fndc-1*(*rny14*) (**f**). **g**, Shown in Fig. 4a. Merged uncropped fluorescence images showing mitochondria (NDUV-2::mNG) and mRb3::FNDC-1 in the −2 oocyte, −1 oocyte and zygote. **h**, Shown in 4f, Supplementary Video 1. Merged uncropped fluorescence images showing mitochondrial network (NDUV-2::mNG) and mRb3::FNDC-1 punctae in the −2 oocyte, −1 oocyte and zygote. Yellow box indicates the region magnified and tracked for time-lapse imaging shown in Fig. 4f. Blue boxes indicate −2 oocytes, violet boxes indicate −1 oocytes, and pink boxes indicate zygotes. Scale bars, 10µm.

**Supplementary Fig. 2:**
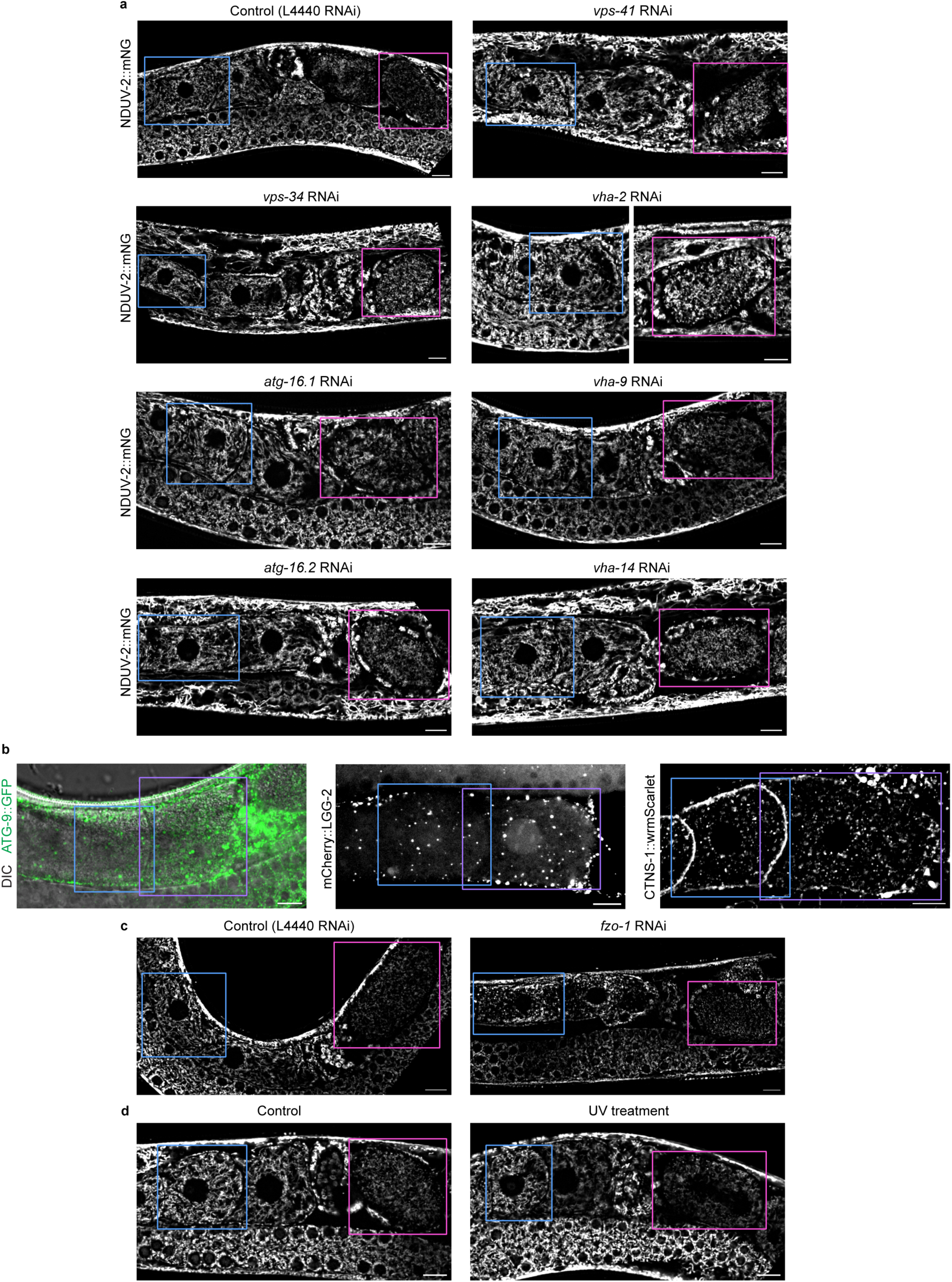
Uncropped confocal fluorescence images from Extended Data figures 2, 3 and 7. **a**, Shown in Extended Data Fig. 2a. Uncropped fluorescence images showing mitochondria (visualized with NDUV-2::mNG) in the −2 oocyte, −1 oocyte and zygote in an empty vector control (L4440) compared to animals treated with RNAi targeting key macroautophagy genes *vps-34*, *atg-16.1*, *atg-16.2*, *vps-41*, *vha-2*, *vha-9* and *vha-14*. **b**, Shown in Extended Data Fig. 2c,d and e. Left to right: Uncropped fluorescence images showing endogenous ATG-9::GFP, germline specific LC3 (mCherry::LGG-2, *pie-1p::mCherry::lgg-2*) and lysosome (visualized with CTNS-1::wrmScarlet) distribution in the −2 oocyte and −1 oocyte. **c**, Shown in Extended Data Fig. 3a. Uncropped fluorescence images showing mitochondria (NDUV-2::mNG) in the −2 oocyte, −1 oocyte and zygote in an empty vector control (L4440) (left) compared to a *fzo-1* RNAi-treated animal (right). **d**, Shown in Extended Data Fig. 7d. Uncropped fluorescence images showing mitochondria (NDUV-2::mNG) in the −2 oocyte, −1 oocyte and zygote in control animal (left) compared to a UVC treated animal (right). Blue boxes indicate −2 oocytes, violet boxes indicate −1 oocytes and pink boxes indicate zygotes. Scale bars, 10µm.

**Supplementary Fig. 3:**
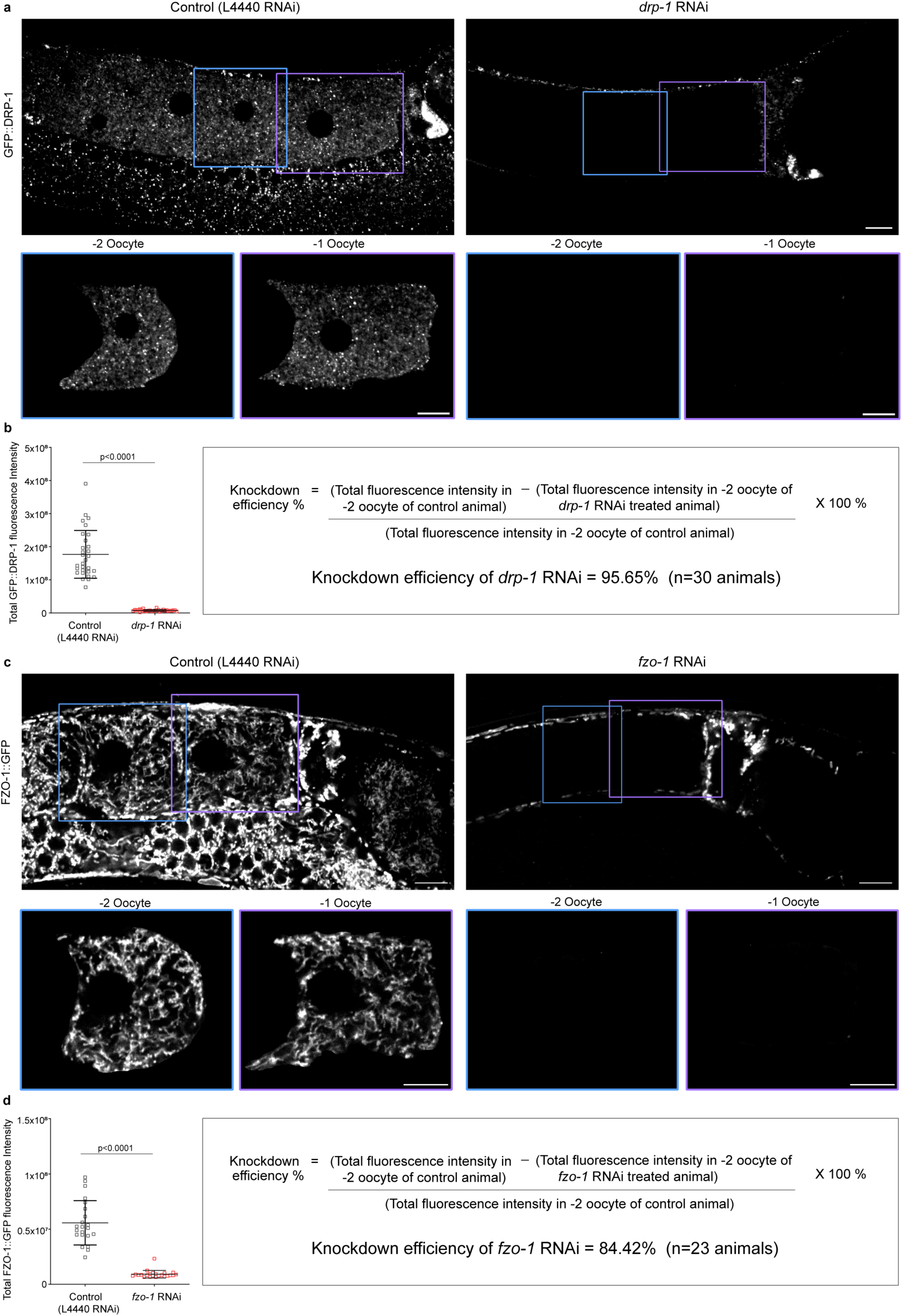
Knockdown efficiency of *drp-1* and *fzo-1* RNAi for experiments in Fig. 2 and Extended Data Fig. 3. **a**, Endogenous GFP::DRP-1 puncta distribution in the germline of an adult empty vector control (L4440) (left) compared to a *drp-1* RNAi-treated animal (right). Blue boxes indicate −2 oocytes and violet boxes −1 oocytes. −2 oocytes and −1 oocytes enclosed in boxed regions are magnified. Scale bars, 10µm. **b**, Left: Total GFP::DRP-1 fluorescence intensity in −2 oocytes compared between empty vector controls (L4440) and *drp-1* RNAi-treated animals. Data represent mean ± s.d. (n = 30 animals per condition) from three biological replicates. P value using two-tailed Student’s *t*-test. Right: Knockdown efficiency calculation for *drp-1* RNAi. **c**, Endogenous FZO-1::GFP (*fzo-1::gfp*) distribution in the germline of an adult empty vector control (L4440) (left) compared to a *fzo-1* RNAi-treated animal (right). Blue boxes indicate −2 oocytes, and violet boxes indicate −1 oocytes. −2 oocytes and −1 oocytes enclosed in boxed regions are magnified. Scale bars, 10µm. **b**, Left: Total FZO-1::GFP fluorescence intensity in −2 oocytes compared between empty vector controls (L4440) and *fzo-1* RNAi-treated animals. Data represent mean ± s.d. (n = 23 animals per condition) from three biological replicates. P value using Mann-Whitney test. Right: Knockdown efficiency calculation for *fzo-1* RNAi.

**Supplementary Table 1.** RNAi screen to identify effectors of change in mitochondrial morphology during the *C. elegans* oocyte-to-zygote transition. The genes targeted by RNAi and effects on mitochondrial morphology observed in oocytes and zygotes of empty vector controls (L4440) and RNAi treated animals across three replicates.

| Mitochondrial fission-fusion RNAi screen |  | Mitochondrial fragmentation during OZT |  |
| --- | --- | --- | --- |
| Gene targeted by RNAi | Number of animals | Oocyte mitochondrial morphology | Zygote mitochondrial morphology |
| L4440 (empty vector control) | 60 | Elongated (n=60/60) | Punctate (n=60/60) |
| *pink-1* | 30 | Elongated (n=27/30) Donut shaped (n=3/30); P=0.1037, ns | Punctate (n=30/30) |
| *drp-1* | 42 | Elongated (n=42/42) | Elongated (n=39/42); P=0.0001 |
| *dyn-1* | 30 | Elongated (n=30/30) | Punctate (n=30/30) |
| *mff-1* | 30 | Elongated (n=30/30) | Punctate (n=30/30) |
| *slc25A46* | 30 | Elongated (n=30/30) | Punctate (n=30/30) |
| *mtp-16* | 30 | Elongated (n=30/30) | Punctate (n=30/30) |
| *fis-1* | 30 | Elongated (n=30/30) | Punctate (n=28/30)<br>Elongated (n=2/30); P=0.1086, ns |
| *fis-2* | 30 | Elongated (n=30/30) | Punctate (n=30/30) |
| *chch-3* | 30 | Elongated (n=30/30) | Punctate (n=30/30) |
| *eat-3* | 30 | Elongated (n=30/30) | Punctate (n=30/30) |
| *mfn-1* | 30 | Elongated (n=30/30) | Punctate (n=30/30) |
| *fzo-1* | 38 | Punctate (n=24/38) Donut + punctate shaped (n=14/30); P=0.0001 | Punctate (n=38/38) |
| *miga-1* | 30 | Elongated (n=30/30) | Punctate (n=30/30) |
| Statistical analysis was conducted using 2x2 contingency table- Fisher's exact test. P values denoted are for comparisons between L4440 control and respective RNAi treated animals. |  |  |  |

**Supplementary Table 2.**
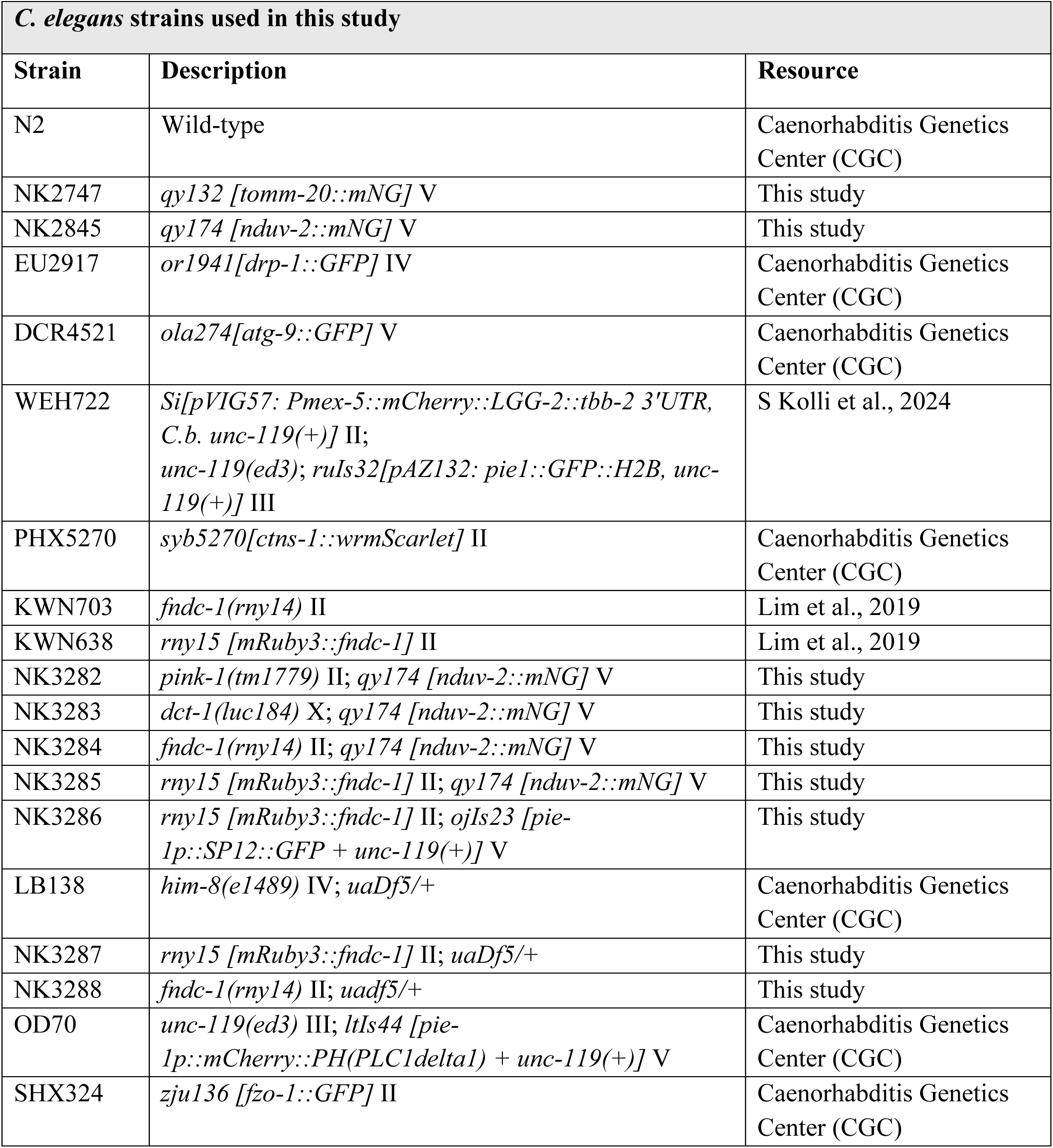
*C. elegans* strains used in the study. Strain name, genotype description, and resource used.

**Supplementary Table 3.**
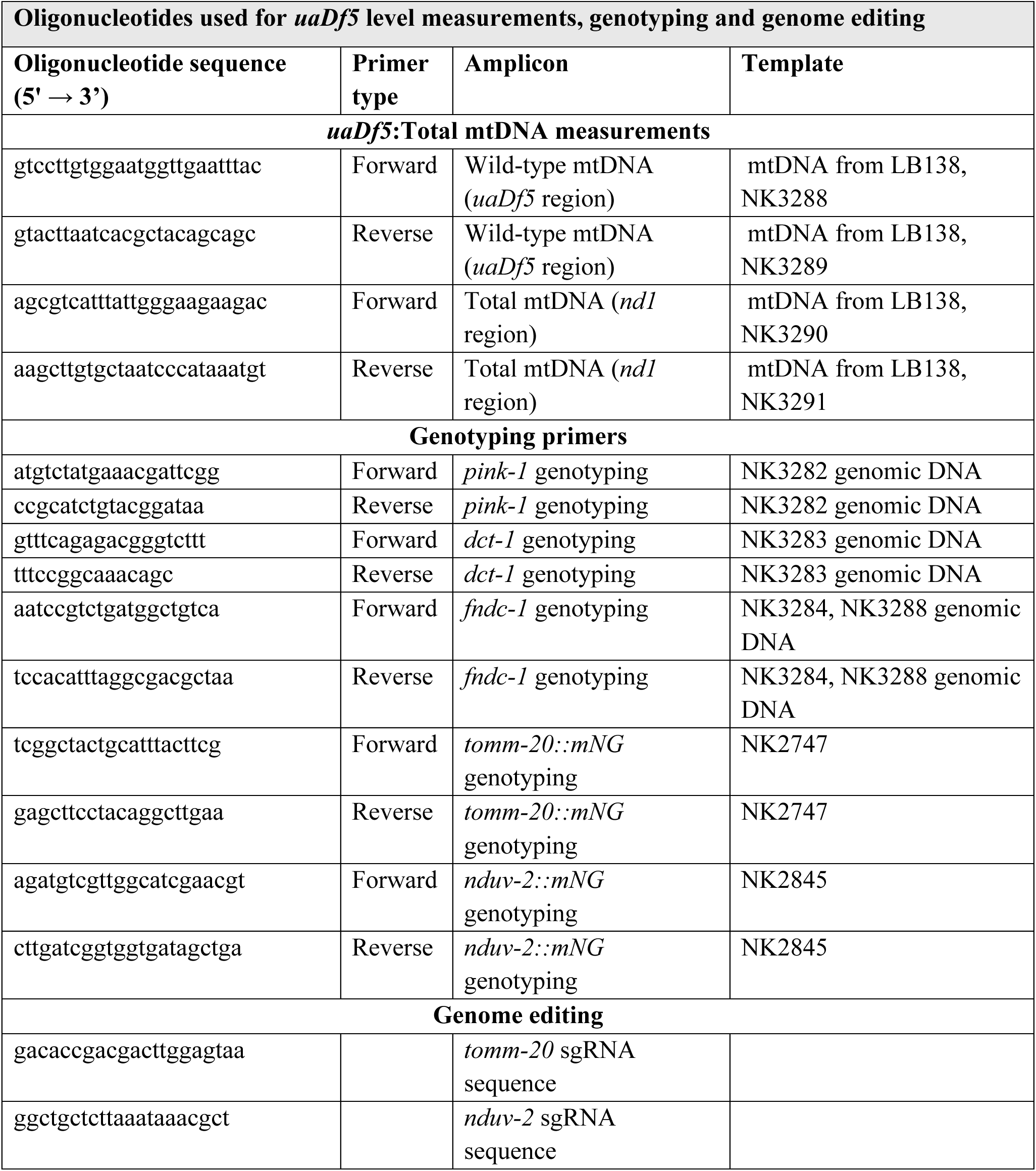
Oligonucleotides used in the study. Oligonucleotides used for *uaDf5* and total mtDNA level measurements, genotyping strains, and genome editing.

